# Multi-Omics analyses reveal dynamic interactions between DNA methylation and transcriptional regulation during black raspberry ripening

**DOI:** 10.1101/2025.11.24.690302

**Authors:** Wei-Hsun Hsieh, Yu-Hung Hung, Jian-Hui Zhang, Han-Yi Chen, Liang-Peng Lin, Meng-Hsun He, Yen-Chng Wang, Yu-Yu Ho, Gina Fernandez, Penelope Perkins-Veazie, Brandon H. Le, Xu Li, Tzung-Fu Hsieh, Jer-Young Lin

## Abstract

We profiled developmental-series transcriptome, methylome, and metabolome profiling to reveal extensive epigenetic reprogramming during black raspberry (*Rubus occidentalis*) fruit ripening. Fruit tissues exhibit globally higher DNA methylation than leaves, particularly in CHG and CHH contexts. Local methylation in all cytosine contexts progressively decreases in promoter regions during ripening, whereas CHG and CHH methylation increase in transposon-rich regions. Two primary methylation transitions—promoter hypomethylation and CHH hypermethylation—govern transcriptional shifts in genes involved in ripening processes. Methyl-binding transcription factors with activation potential likely promote CHH hypermethylation-linked transcriptional activation. Multi-omics integration revealed coordinated anthocyanin accumulation parallels expression of biosynthetic and regulatory genes within coherent networks. Elevated non-CG methylation in heterochromatin coincides with increased transcription of histone variants mediating chromatin compaction, suggesting chromatin remodelers fine-tune accessibility for methyltransferases. Our study highlights both genome-wide and locus-specific epigenetic reprogramming and demonstrates a coordinated interplay between DNA methylation and transcriptional regulation during black raspberry fruit ripening.

## Introduction

Fleshy fruit ripening is characterized by physiological modifications in color, texture, sugar content, aroma, and flavor, enhancing their attractiveness to seed-dispersal vectors. This developmental transition is tightly controlled in a tissue- and stage-specific manner, and misregulation results in deleterious consequences^1^. Fruits have traditionally been categorized into climacteric and non-climacteric groups. The hormone responses, genetic factors and molecular mechanisms underlying the ripening processes in climacteric fruits, such as tomato, are relatively well documented^1^. Compared with climacteric fruits, the ripening process in non-climacteric fruits such as strawberry, grape, and citrus remains poorly understood; however, recent studies have begun to uncover the molecular mechanisms underlying this process. Non-climacteric fruits, such as black raspberry, lack the characteristic respiratory burst and surge in ethylene production observed in climacteric species. Emerging research suggests that ripening in non-climacteric fruits is associated with multiple hormones, such as ABA, GA, and auxin^1^. In addition to hormonal regulation, the fruit ripening process is also coordinated by transcription factors (TFs) and epigenetic modifications^2,3^. In climacteric tomato, the TF CNR gene is silenced in the colorless non-ripening (Cnr) mutant due to promoter hypermethylation, arresting fruit development^4^. Another important regulator, RIPENING INHIBITOR (RIN), a MADS-box TF, activates numerous ripening-related genes marked by promoter hypomethylation^5^. Disruption of the DNA demethylase SlDML2 in tomato delays ripening^6,7^, suggesting that active DNA demethylation underlying this developmental program. Additionally, alterations in DNA methylation at MYB loci have been associated with changes in anthocyanin biosynthesis in climacteric apple and climacteric pear^8,9^. These studies demonstrated that changes in DNA methylation play a pivotal role in fruit ripening.

Shifts in DNA methylation patterns accompany fruit ripening in both climacteric and non-climacteric species. For instance, global hypermethylation has been observed in both climacteric fruits (pear, apple) and non-climacteric fruits (orange, grape), while hypomethylation also occurs in climacteric tomato as well as in non-climacteric strawberry^10–13^. These patterns suggest that the DNA methylation change during fruit ripening is shaped by fruit-specific program rather than climacteric classification. Moreover, the molecular basis driving global DNA methylation changes differs across species. The global hypomethylation in non-climacteric strawberry is linked to suppression of the RNA-dependent DNA methylation (RdDM) pathway genes, whereas in climacteric tomato, it is driven by enhanced expression of the DNA demethylase^5,14^. Meanwhile, hypermethylation in both climacteric apple and non-climacteric orange results from downregulation of DNA demethylase genes^11,13^. These distinct methylation pathways support the idea that fruit-specific DNA methylation changes are driven by species-specific mechanisms.

Black raspberry (*Rubus occidentalis*) fruits have high levels of anthocyanins, contributing to its exceptional antioxidant capacity and associated health benefits^15^. Each fruit is an aggregate of many individual drupelets. Each drupelet is anatomically analogous to a cherry, featuring a hard endocarpic seed (pyrene) surrounded by a fleshy mesocarp and an outer exocarp consisting of a thin skin (http://www.efloras.org/). After fertilization, fruit development progresses from an early stage of intense cell division to reduced proliferation as the embryo and seed coat mature, culminating in rapid fruit expansion driven by extensive cell enlargement (Fig. 1A). Ripening studies in this species may provide insights for extending shelf life and enhancing nutritional quality. Black raspberry fruit ripening involves complex physiological and biochemical changes that make the fruit soft, flavorful, and rich in nutrients. Despite its agricultural and nutritional importance, the molecular basis of this ripening process—especially the role of DNA methylation in regulating gene expression—remains largely unexplored.

**Fig. 1.**
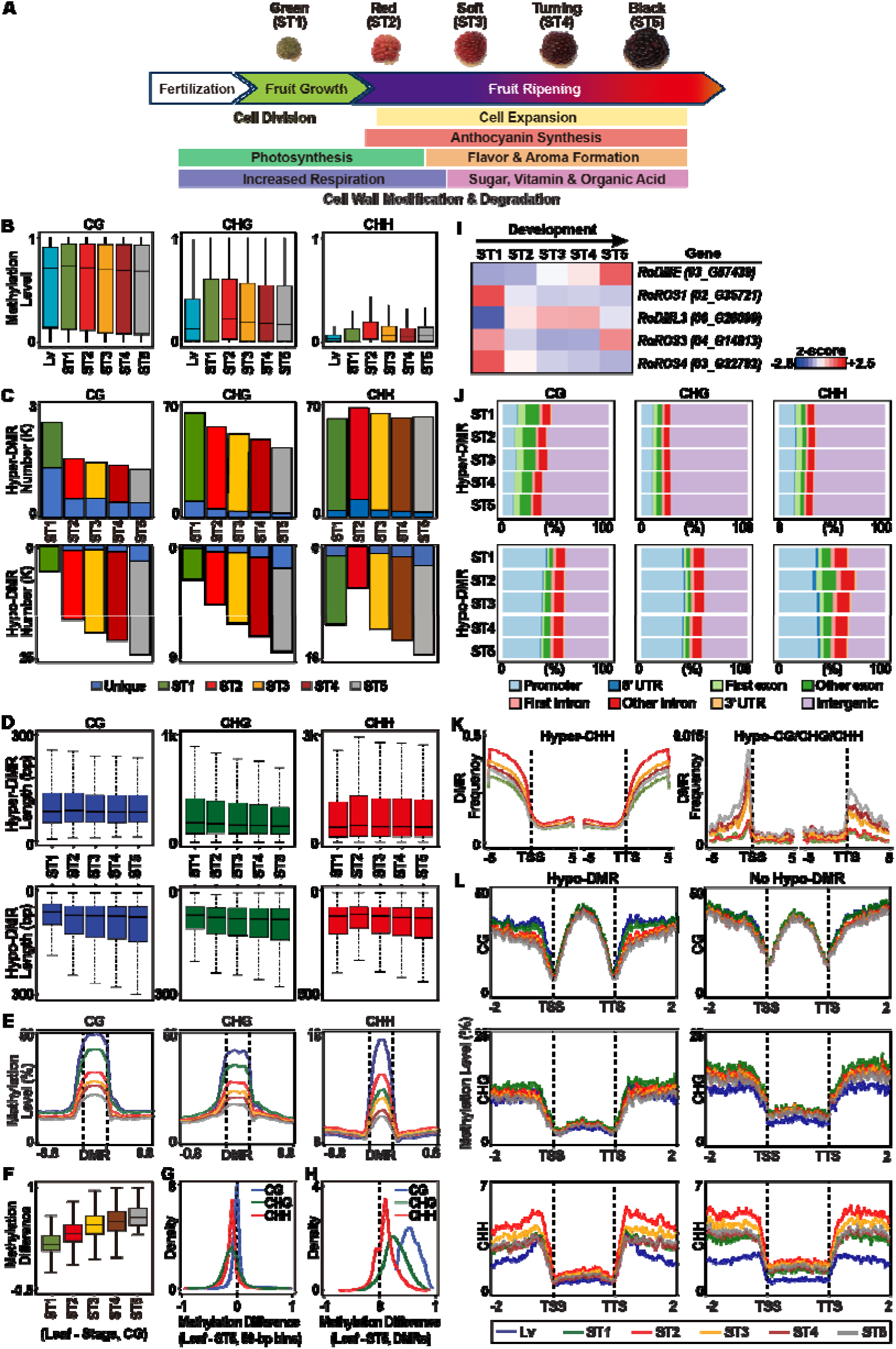
Schematic representation of black raspberry fruit developmental stages and associated DNA methylation features during ripening. **A** Black raspberry fruit ripening stages and major developmental events. **B** DNA methylation levels at five fruit ripening stages and in leaf. Methylation levels are shown as box plots representing 500-kb windows across the genome. **C** Number of hyper- and hypo-DMRs at each fruit ripening stage compared to leaf. **D** Lengths of differentially methylated regions (DMRs), including both hypo- and hyper-DMRs, during black raspberry fruit ripening. Leaf tissue was used as the reference for comparing DMRs at each stage. **E** DNA methylation patterns of gene-proximal CG, CHG, and CHH hypo-DMRs, derived from ST5-to-leaf comparison, across black raspberry fruit ripening, including corresponding levels in leaves. **F** DNA methylation levels of CG hypo-DMRs identified in the black stage compared to leaf shown across all ripening stages. **G** Genome-wide DNA methylation differences between the black stage (ST5) and leaf, calculated using 50-bp tiling windows. **H** DNA methylation differences between the black stage and leaf, based on CG hypo-DMRs identified in the black stage relative to leaf. **I** Temporal expression patterns of DNA demethylase genes across ripening stages represented by heatmap. **J** Genomic localizations of DMRs during black raspberry fruit ripening. K DMR frequency in genic regions. **L** DNA methylation levels across genic regions in genes with or without CG, CHG, and CHH hypo-DMRs.

We profiled developmental-series of the transcriptome, methylome, and targeted metabolome to characterize the molecular reprogramming underlying black raspberry (*Rubus occidentalis*) fruit ripening. Global DNA methylation in the fruit is reprogrammed, with overall levels significantly higher than in the leaf, particularly in CHG and CHH contexts. Local hypomethylation in CG, CHG, and CHH, predominantly enriched in promoter regions, show progressive demethylation across ripening, evident in both the increasing number of demethylated loci and the decreasing methylation levels. By contrast, local hypermethylation in CHG and CHH contexts are predominantly located in intergenic and transposon-rich heterochromatic regions. Transcriptional reprogramming during fruit development is associated with two predominant types of local methylation changes: promoter-associated CG/CHG/CHH hypomethylation and CHH hypermethylation, both of which correlate with altered expression of genes involved in fruit physiological process. The developmentally regulated expression of methyl-binding transcription factors with transcriptional activation potential likely contributes to the elevated CHH methylation associated with increased transcriptional activity. Multi-omics integration revealed correlated accumulation of anthocyanin species with the expression of specific biosynthetic and regulatory genes interconnected within defined gene regulatory networks. Elevated non-CG methylation in heterochromatin, relative to the leaf, coincides with increased transcription of histone variants involved in forming compact heterochromatin that is inaccessible to DNA methyltransferases. This paradox may be partially explained by the diverse chromatin remodelers active during ripening, which modulate chromatin accessibility for methyltransferases. Our study reveals genetic and epigenetic programs that orchestrate fruit maturation and anthocyanin biosynthesis in black raspberry fruit.

## Results

### Generation of single-base resolution black raspberry fruit methylomes throughout development and ripening

We generated single-base resolution DNA methylation profiles of black raspberry fruit across five developmental and ripening stages using bisulfite sequencing (BS-Seq), capturing the full progression to compare the methylation landscape of reproductive tissues with that of the somatic tissue (leaf). The stages were designated as the (i) green (stage 1, ST1), (ii) red (stage 2, ST2), (iii) soft (stage 3, ST3), (iv) turning (stage 4, ST4) and the full ripening black (stage 5, ST5) stages (Fig. 1A). In total, we generated ∼1.06 billion Illumina BS-Seq reads from all fruit stages and leaf, obtaining in each case 16.1–43.8× coverage of each strand in the 300-Mb black raspberry genome (Supplementary Table 1A). We assayed 97.1–99.5 million cytosines, representing 89.0–91.2% of all cytosines in the genome. Across the fruit samples, the average methylation levels were ∼55% (CG), ∼24% (CHG), and ∼5% (CHH, H = C, A or T), corresponding to ∼64%, ∼39%, and ∼20% of cytosines methylated in each context (Supplementary Table 1B). The unmethylated λ genome (added to our samples as an internal control) shows >99% C-to-T conversion, indicating high bisulfite conversion efficiency in BS treatment (Supplementary Table 1C). The BS-Seq data from biological replicates of fruits were in excellent agreement with each other (correlation coefficients > 0.99) (Supplementary Figure 1). Taken together, these results indicate that our datasets represent unbiased, deep representation, and highly reproducible profiles of a temporal DNA methylation changes across the black raspberry genome throughout fruit ripening.

### Global DNA methylation levels in fruit ripening are higher than in leaf

To determine global DNA methylation levels, the number and fraction of methylated cytosines in 50-bp intervals were counted on each strand across the entire methylome. Relative to leaf, the fruit genome undergoes genome-wide DNA methylation changes, including a slight decrease in CG methylation and substantial increases in CHG and CHH methylation (Fig. 1B and Supplementary Figure 2A). Compared with the green (ST1), CG methylation is slightly reduced in later stages, whereas CHG methylation declines and CHH methylation increases (Fig. 1B and Supplementary Figure. 2B); these analysis results are similar to the whole genome average methylation levels (Supplementary Figure 2C). The distribution of methylated cytosine sites in the fruit genome uncovers that symmetric methylation (CG and CHG) gradually declines from ST1 to ST5, whereas CHH methylation peaks at ST2 and then decreases during later stages (Supplementary Figure 2C, 2D). Together, these results showed that the genome-wide methylation patterns are reprogrammed during fruit development and were elevated in the fruit compared with the leaves, suggesting distinct methylation dynamics in the reproductive and somatic tissues.

### Local DNA methylation patterns are changed across fruit ripening

Next, we investigated whether there were changes in local DNA methylation patterns (differentially methylated regions, DMRs) across fruit ripening. We applied CGmapTools to identify DMRs with the cutoff *p*-value < 0.05, CG > 20%, CHG > 10% and CHH > 8%^16^. To precisely and comprehensively inspect the occurrence of DMRs in fruit, DNA methylation levels in leaf were used as baseline references for comparison with those in each fruit ripening stage. We found that there were more hypo-DMRs than hyper-DMRs in the CG context, whereas in the non-CG contexts, hyper-DMRs outnumbered hypo-DMRs (Fig. 1C and Supplementary Figure 3). We further found that the number of hyper-DMRs decreased in the CG and CHG contexts, while the number of hypo-DMRs increased during fruit ripening (Fig. 1C and Supplementary Figure 3). Remarkably, most DMRs occurred in the same loci with very few being unique in pair-wise comparison. The lengths of hypo-DMRs in all three cytosine contexts increased as the fruit developed, while those of the hyper-DMRs remained the same or slightly decreased (Fig. 1D). The increase in length and the limited number of unique new hypo-DMRs in successive stages indicated that most demethylations occurred on the same loci, which may suggest that the methylation levels in the DMRs continue to decrease and that the demethylated regions continue to expand.

### Progressive active DNA demethylation at specific loci during fruit ripening

We examined DNA methylation levels within the ST5-vs-leaf hypo-DMRs across all developmental stages and found a continued decline in methylation in each cytosine context as ripening progressed, indicating that these hypo-DMRs progressively lose DNA methylation during fruit ripening (Fig. 1E). Consequently, the CG methylation difference continued to increase in these same CG hypo-DMRs as fruit ripening progressed (Fig. 1F). We further examined the distribution patterns of CHG and CHH methylation difference within these CG hypo-DMRs. As a control, we inspected the methylation difference genome-wide with respect to 50-bp bins. We found that the CHG and CHH methylation levels of the 50-bp bins were generally higher in ST5 relative to leaves (density peaks shifted to the negative side), while CG methylation levels were similar in both tissues (Fig. 1G).

However, the CHG and CHH methylation levels within the ST5 CG hypo-DMRs relative to leaf were also predominantly lower than in the leaf (Fig. 1H), indicating that these ST5 CG hypo-DMRs were demethylated in all cytosine contexts. As cell division ceases following early fruit ripening, the extensive DNA demethylation detected is most likely attributed to active DNA demethylation rather than passive loss through replication. Supproting this model, various DNA demethylases were active at different ripening stages: *RoROS1,3,4* were highly active at ST1, whereas *RoDME, RoDML3* and *RoROS3* were active at ST3 to ST5, suggesting dynamic and coordinated active DNA demethylation throughout the fruit ripening (Fig. 1I and Supplementary Table 2A). Together, the results indicated that compared to the leaf, black raspberry fruit ripening involves locus-specific reductions in CG, CHG and CHH methylation, accompanied by widespread increases in CHG and CHH methylation genome-wide, highlighting a concerted epigenomic remodeling.

### Hypo-DMRs are located primarily in promoter and gene body regions

We examined the genomic locations of the DMRs and found that hyper- and hypo-DMRs are preferentially distributed across different genomic regions. Most hypo-DMRs in all three cytosine contexts were located in genic regions (Fig. 1J), out of which more than half were in promoters; by contrast, most hyper-DMRs were found in intergenic regions (Fig. 1J). Additionally, these non-CG hyper-DMRs were co-localized with chromosomal regions enriched for LTR retrotransposons (Supplementary Figure 4). Consistent with the decrease in non-CG methylation levels in the CG hypo-DMRs, many non-CG hypo-DMRs overlapped with CG hypo-DMRs (Supplementary Figure 5). The frequency of CG/CHG/CHH hypo-DMRs was greater in regions upstream and proximal to the transcription start site (TSS) than in regions downstream of the transcription termination site (TTS), and DMRs in both ends diminished progressively in the distal regions (Fig. 1K). By contrast, CHH hyper-DMRs accumulated near both the TSS and TTS and increased successively in the distal regions (Fig. 1K). These distinct spatial patterns across genic regions indicate that various epigenetic modifications may shape regulatory landscapes flanking genic regions in a context-dependent manner. Interestingly, genes containing CG/CHG/CHH hypo-DMRs exhibited lower DNA methylation levels around both the TSS and TTS compared with genes lacking these DMRs (Fig. 1L), suggesting that distinct molecular mechanisms may underlie the establishment of DNA methylation landscapes in genes with hypo-DMRs relative to those without them.

Overall, DNA methylation landscapes differed substantially between the leaf and fruit. Early in fruit development, DNA methylation declined in specific promoters and genic regions, presumably mediated by the DNA demethylases active in early ripening (Supplementary Figure 6). As ripening progressed, active demethylation continued and expanded in the same loci, as well as at newly targeted sites via additional stage-preferred demethylases (Supplementary Figure 6). By contrast, relative to leaf, non-CG methylation increased mainly in non-genic regions, particularly within repetitive sequences (Supplementary Figure 4). Our results, therefore, suggest that, black raspberry fruit ripening involves multiple mechanisms of DNA methylation reprogramming.

### Extensive transcriptomic reprogramming occurs at the soft fruit stage

To understand the genome-wide transcript changes that drive black raspberry fruit ripening, we analyzed genome-wide gene expression patterns across five fruit stages (Fig. 1A). We inspected the correlations of whole genome transcripts between bio-replicates using principal component analysis (PCA) to demonstrate that bio-replicates are closely correlated with each other and that transcripts between different stages are distinct, indicating that distinct sets of genes are sequentially activated or repressed to coordinate the progression of fruit ripening (Supplementary Figure 7). We applied unsupervised hierarchical clustering analysis to inspect the relationships of whole genome transcripts between fruit ripening stages (Fig. 2A). The transcriptomes across ripening stages can be divided into two groups: (i) an early group from green to red (ST1 and ST2), and (ii) a late group from soft to black (ST3 to ST5). Significantly, soft (ST3) is distal from the other two stages (ST4 and ST5) in the late group, representing a transition marked by a distinct mRNA population. To uncover the genes that are active or repressed in each stage compared to other stages, we applied DESeq2 (FDR < 0.05) with more than a two-fold change to identify differentially expressed genes (DEGs). During black raspberry fruit ripening, 14,544 genes (representing 43.7% of the annotated genome) are differentially expressed, with a greater number of genes down-regulated than upregulated (Fig. 2B). In sequential stage-to-stage comparisons, more DEGs were identified during the early transition stages (from ST1 to ST2 and ST2 to ST3) than in the later transition stages (ST3 to ST4 and ST4 to ST5), indicating that large-scale transcriptomic reprogramming occurs primarily during early fruit development. Notably, the transition from ST2 to ST3 was marked by the most pronounced transcriptional shift among all successive-stage comparisons, with 2,191 genes upregulated and 3,914 genes downregulated relative to ST2. To uncover the biological processes regulated by transcriptional programs during fruit ripening, we performed Gene Ontology (GO) analysis on DEGs between consecutive developmental stages (Supplementary Figure 8 and Supplementary Table 3). Genes upregulated at the soft stage (ST3) were found to be enriched in biological functions including fatty acid biosynthesis, glycolysis, flavonoid biosynthesis and ABA signaling, while downregulated genes were associated with processes such as photorespiration and mitochondrial organization. These functions reflect many physiological signatures of the fruit ripening processes. Together, these transcriptome analyses revealed two transcriptionally distinct phases during black raspberry fruit ripening, suggesting extensive transcriptional reprogramming.

**Fig. 2.**
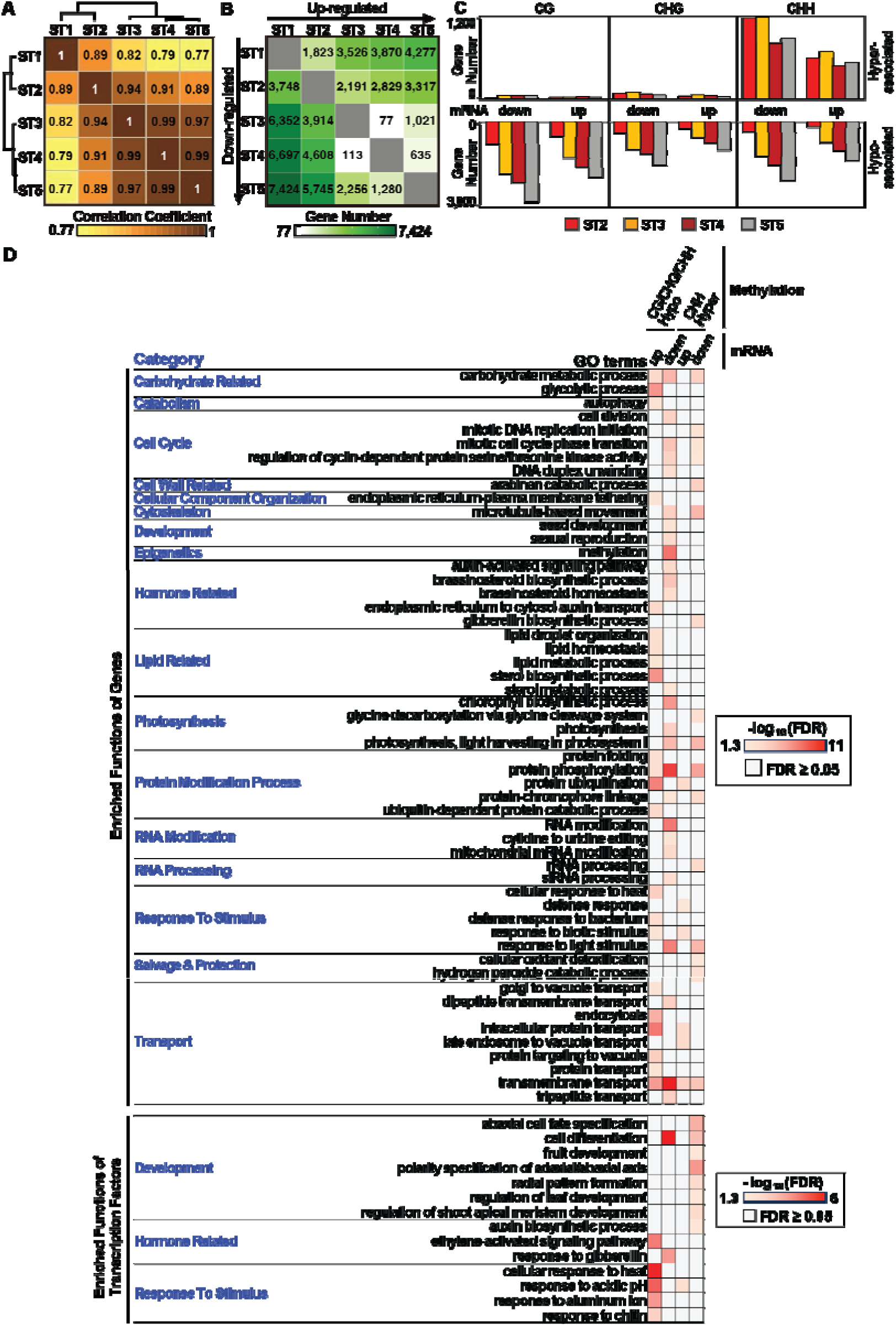
Gene expression dynamics and the overlap of differentially expressed genes (DEGs) and differentially methylated genes (DMGs), along with their enriched functional categories during black raspberry fruit ripening. **A** Correlation coefficient analysis of RNA populations between stages was performed, and the resulting coefficients were hierarchically clustered to illustrate the association of ripening stages. **B** Differentially expressed genes in a pair-wise comparison. The numbers in the matrix represent the DEG numbers. The left side of the diagonal represents genes down-regulated in one stage on the Y-axis compared to the X-axis. The right side of the diagonal represents genes up-regulated in one stage on the X-axis compared to the Y-axis. **C** Bar plot summarizing genes that are both DEGs and DMGs, using ST1 as the reference. mRNA down: genes downregulated relative to ST1; mRNA up: genes upregulated relative to ST1. **D** Enriched Gene Ontology (GO) biological process terms for DEGs and differentially expressed transcription factors associated with CG, CHG, and CHH hypo-DMRs, as well as CHH hyper-DMR. mRNA down: downregulated genes; mRNA up: upregulated genes.

### The activities of a subset of genes are associated with DNA methylation changes

To examine the relationship between local DNA methylation changes and gene activities, we identified differentially methylated genes (DMGs) as those harboring DMRs within the promoter (1.5 kb upstream of the TSS) and/or gene body. By intersecting DMGs with DEGs, we identified a subset of genes whose expression changes were associated with alterations in local DNA methylation (Fig. 2C and Supplementary Figure 9). DEGs associated with hypo-DMRs in all cytosine contexts or with CHH hyper-DMRs far exceeded those associated with CG or CHG hyper-DMRs, suggesting CG/CHG/CHH hypomethylation or CHH hypermethylation may dynamically contribute to the transcriptional regulation in black raspberry. We examined the major biological functions of the DEGs-associated DMGs (Fig. 2D and Supplementary Table 4). Among the upregulated genes with CG/CHG/CHH hypo-DMRs, enriched functions included glycolysis, lipid metabolism and transport. By contrast, downregulated genes with CG/CHG/CHH hypo-DMRs were primarily associated with cell cycle regulation, photosynthesis (e.g., *CRD1* and *DXS1*) and hormone-related processes such as the brassinosteroids, GA and auxin (e.g., *ARF1*, *LRP1* and *PILS5*) (Supplementary Figure 10). Additionally, both up and downregulated DEGs linked to hypo-CG/CHG/CHH DMRs or hyper-CHH DMRs were enriched for overlapping functions, including carbohydrate metabolism (e.g., *PGF10*, *6PGD* and *BGLU17*), protein phosphorylation (e.g., *ABC1-like*, *SERK1* and *Haspin*) and transport (e.g., *ABCG40*, *DTX44*) (Supplementary Figure 10). Interestingly, downregulated DEGs associated with CHH hyper-DMRs demonstrated a broader spectrum of functional categories than their upregulated counterparts, suggesting that CHH hypermethylation can influence transcription bidirectionally, with repression as its dominant effect. We further analyzed differentially expressed TF genes associated with DNA methylation changes (Fig. 2D and Supplementary Table 5), revealing that the major functions are related to development, response to stimuli, or hormone-related processes, such as auxin, ethylene, and GA. Given that ripening-related TFs are associated with DNA methylation changes in tomato^5^, we investigated whether ripening-related TFs show similar patterns in black raspberry fruit. Key regulators such as NOR, CNR, and TDR4 exhibited expression changes linked to local DNA methylation alterations (Supplementary Figure 10). Together, these results show that DNA methylation changes are associated with transcriptional reprogramming of genes enriched with functions involved in black raspberry fruit ripening.

### Epigenetic regulators promoting methylated gene expression and DNA methylation are regulated at the transcriptional level

Although DNA methylation is traditionally associated with transcriptional repression, our results demonstrate that elevated CHH methylation can also coincide with gene activation enriched in specific functions (Fig. 2D). This pattern aligns with recent studies showing that specific methyl-binding TFs and proteins (e.g., LEC2, DNAJ2, SUVH1, SUVH3, and MBD9) are capable of binding methylated DNA and promoting gene expression in specific loci^17–20^. Conversely, other methyl-binding TFs and proteins, such as members of the TF REM family, VIM1, and SUVH4/5/6, can recruit DNA methylation machinery (e.g., RdDM pathway complex) to establish locus-specific methylation^17,21^. Transcriptional profiling of methyl-binding TFs and proteins during fruit ripening revealed two expression patterns (Fig. 3, Supplementary Table 2B): an early group (e.g., RoLEC2, RoSUVH1/3, and REM TF family members) and a late group (e.g., RoMBD9 and RoDNAJ2). These results suggested that sequence-specific TFs and methyl-binding proteins direct transcriptional activation or DNA methylation in a locus- and stage-specific manner during fruit ripening.

**Fig. 3.**
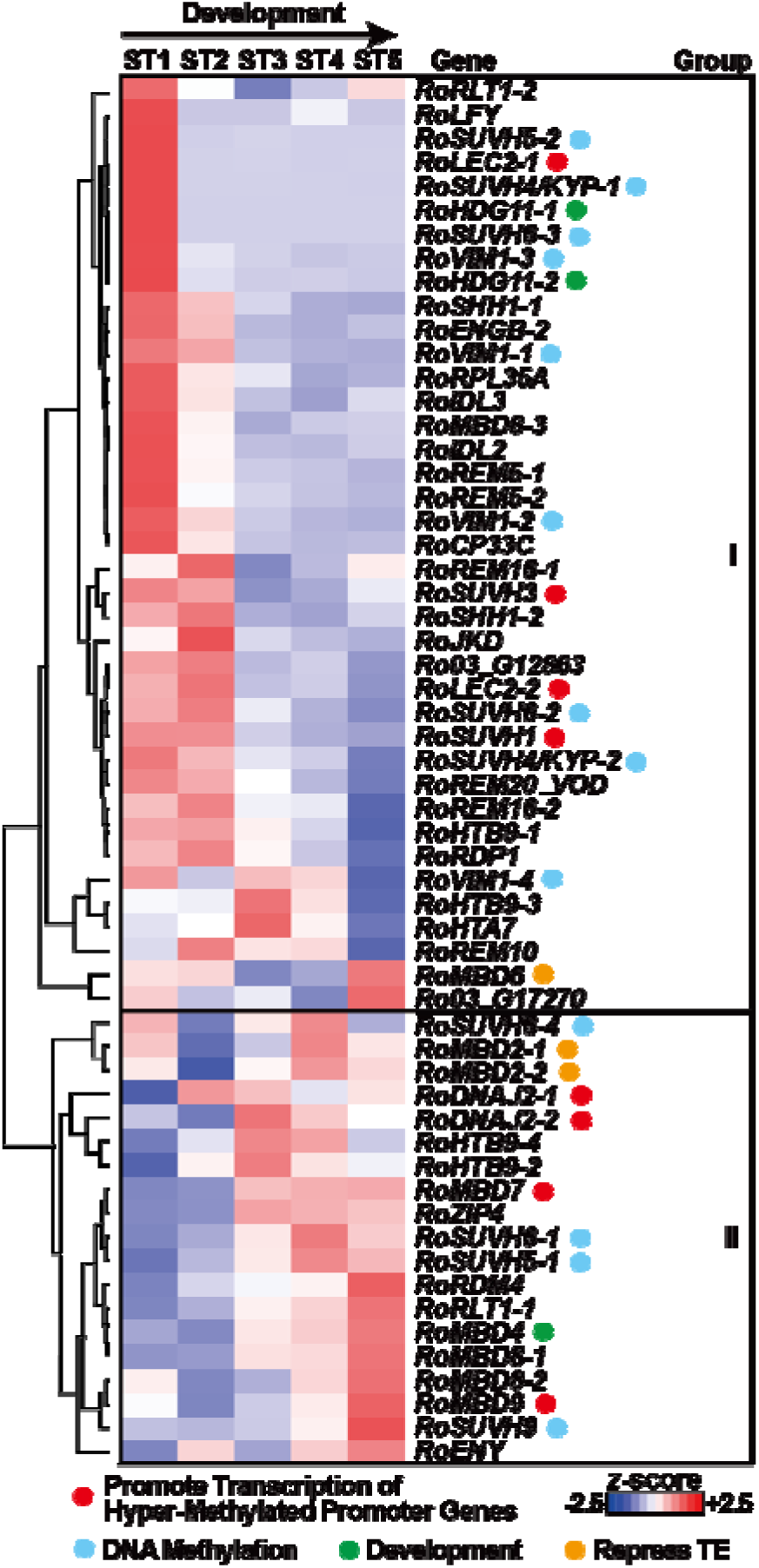
Expression profiles of genes encoding methyl-binding transcription factors and proteins. Unsupervised hierarchical clustering reveals two major transcript groups across black raspberry ripening stages. Genes marked with different colored circles represent distinct biological functions.

### Anthocyanin compounds are predominantly synthesized from the red stage through the black stage

To monitor the temporal accumulation patterns of anthocyanin species, we profiled metabolomes throughout black raspberry fruit ripening. We detected 89 metabolites, including seven anthocyanin species known to be produced in black raspberry fruit^22^ (Supplementary Table 6). We used unsupervised hierarchical clustering to examine the relative abundance of detected anthocyanins across ripening stages, identifying two main groups: (i) those that increase substantially at red stage (ST2; cyanidin 3-glucoside, cyanidin 3-sambubioside, cyanidin 3-xylosylrutinoside, and cyanidin 3-rutinoside), and (ii) those that rise at soft stage (ST3; pelargonidin 3-glucoside, pelargonidin 3-rutinoside, and peonidin 3-rutinoside) (Fig. 4A). Together, these results suggest that anthocyanin biosynthesis is developmentally regulated and primarily initiated at the red or soft stage.

**Fig. 4.**
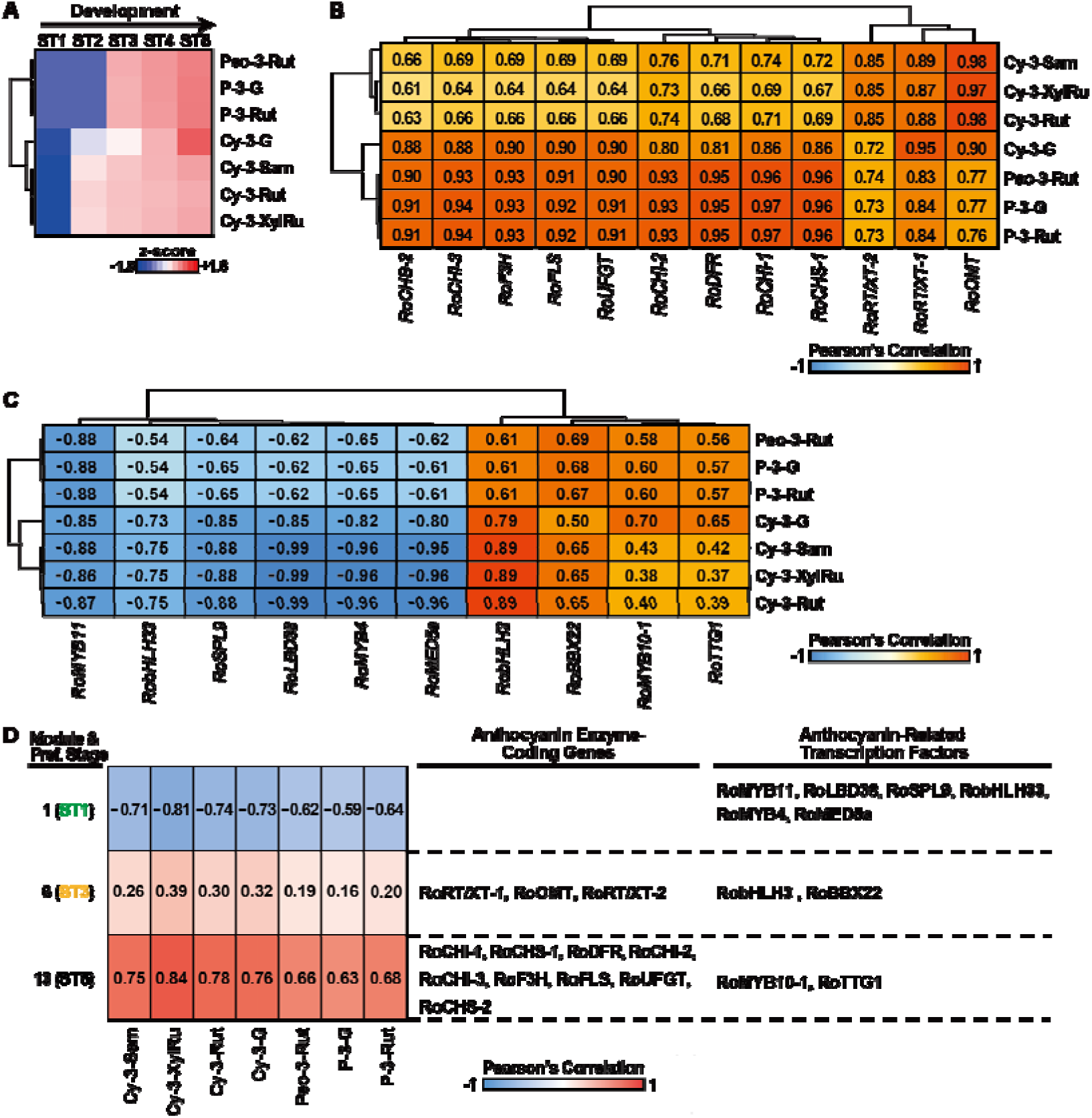
Metabolome analysis of anthocyanin accumulation. **A** Relative levels of anthocyanin compounds clustered by unsupervised hierarchical clustering across ripening stages. **B** Pearson correlation coefficients between transcript levels of 12 enzyme-coding genes in the anthocyanin biosynthetic pathway and the relative abundance of seven anthocyanin compounds. **C** Pearson correlation coefficients between transcript levels of transcription factors regulating anthocyanin biosynthetic genes and the abundance of seven anthocyanin compounds. **D** Module–trait heatmap showing the correlation between gene expression modules and anthocyanin content. Color gradients indicate the strength and direction of the correlations. Cy-3-Sam, cyanidin 3-sambubioside; Cy-3-XylRu, cyanidin 3-xylosylrutinoside; Cy-3-Rut, cyanidin 3-rutinoside; Cy-3-G, cyanidin 3-glucoside; Peo-3-Rut, peonidin 3-rutinoside; P-3-G, pelargonidin 3-glucoside; P-3-Rut, pelargonidin 3-rutinoside.

### Anthocyanin accumulation correlates with the expression of biosynthetic genes and their regulatory transcription factors

To understand how the anthocyanin accumulation is modulated at the transcriptional level during black raspberry ripening, we examined transcript levels of anthocyanin biosynthetic genes in parallel with the relative abundance of seven detected anthocyanin compounds across stages (Fig. 4B and Supplementary Figure 11). The putative enzymes involved in the metabolic pathways of these seven anthocyanins were deduced based on similar pathways in other plant species^23^ (Supplementary Information). Using a correlation coefficient cutoff of *r* ≥ 0.8, we identified 12 transcripts of enzyme-coding genes that are strongly associated with anthocyanin levels: (i) early biosynthetic genes (EBGs) involved in anthocyanin biosynthetic pathway, such as chalcone synthase (CHS), chalcone isomerase (CHI), and flavanone 3-hydroxylase (F3H, key enzyme in the anthocyanin pathway), and (ii) late biosynthetic genes (LBGs), such as flavonoid 3-O-glucosyltransferase (UFGT) and O-methyltransferase (OMT) (Fig. 4B and Supplementary Figure 11). We further investigated whether the expression patterns of the TFs known to regulate anthocyanin biosynthetic genes are also correlated with anthocyanin accumulation. Using the same approach, we identified several TF transcripts that are either strongly positively or negatively correlated (*r* ≥ 0.8 or *r* ≤ –0.8) with anthocyanin levels (Fig. 4C and Supplementary Figure 11). For example, RobHLH3, a component of the MBW complex that targets LBGs^24^, was strongly positively correlated with anthocyanin content, while RoSPL9, a negative regulator of anthocyanin production that destabilizes the MBW complex^25^, exhibited a strong negative correlation with anthocyanin levels (Fig. 4C and Supplementary Figure 11). Finally, we examined the DNA methylation patterns of anthocyanin pathway genes and found that several of them, including *RoFLS* and *RoCHS-1*, show methylation changes associated with altered transcript levels. A similar pattern was observed in the TFs regulating these pathways, such as *RoMYB10-1* and *RoMYB11* (Supplementary Figure 10). Together, the results suggest that anthocyanin accumulation and the expression of anthocyanin biosynthetic genes are coordinated, and their transcriptional regulation may be involved in modulating anthocyanin levels during black raspberry fruit ripening.

### Interconnecting anthocyanin biosynthetic genes and their transcription factors within gene networks coordinating anthocyanin accumulation

To characterize the gene regulatory events programing the black raspberry fruit ripening and governing anthocyanin accumulation, we first inferred gene regulatory network using weighted gene coexpression network analysis (WGCNA) of highly regulated genes (DESeq2 FDR < 0.05 and >2-fold change) (Supplementary Information). A total of 14,457 genes were organized into 14 gene regulatory modules. To understand the temporal relationships between gene modules and ripening stages, we calculated correlation coefficients and performed hierarchical clustering on the resulting r values (Supplementary Figure 12). This revealed stage-specific gene modules, such as module 10, which were strongly associated with the black stage. To further uncover the genetic architecture of anthocyanin biosynthesis, we integrated anthocyanin accumulation patterns with inferred gene networks to assess the relationships between gene modules and anthocyanin contents. We found that the seven anthocyanins are highly associated with module 6 and 13 (Fig. 4D). Specifically, the identified anthocyanin biosynthetic genes (Fig. 4B) were enriched in (i) module 6 (Fig. 4D), associated with ST3 (the soft stage; Supplementary Figure 12), and in (ii) module 13 linked with ST5 (the black stage; Supplementary Figure 12). These two modules contained four TFs (RobHLH3, RoBBX22, RoMYB10-1 and RoTTG1) (Fig. 4D) whose expression levels were positively correlated with anthocyanin accumulation (Fig. 4C). Therefore, the consistent positive associations observed between mRNA expression and anthocyanin levels (Fig. 4B, C), as well as between modules and anthocyanin accumulation (Fig. 4D), suggest that these modules may act as positive regulatory hubs for fruit pigmentation. By contrast, TFs negatively correlated with anthocyanin content—such as the CHS repressor RoMYB4 (Fig. 4C)—were enriched in module 1. This module operates during ST1 (the green stage; Supplementary Figure 12), characterized by low anthocyanin levels (Fig. 4A). Its strong negative correlation with anthocyanin accumulation (Fig. 4D) indicates a repressive role in early-stage of anthocyanin biosynthesis. Notably, MED5a, a Mediator complex subunit known to be involved in transcriptional repression of the phenylpropanoid metabolism in *Arabidopsis* ^26^, is a constituent of this negative correlating module 1 (Fig. 4D). Its presence reinforces the view that module 1 likely mediates repression of anthocyanin biosynthesis in black raspberry, attesting to the robustness and biological relevance of our gene regulatory network analysis. Together, the results suggested that a coordinated gene regulatory architecture orchestrates the distinct anthocyanin accumulation profiles during black raspberry fruit ripening.

### Non-CG methylation levels in heterochromatin in the fruit genome are higher than in leaf

In our DMR analysis, we uncovered that hyper-DMRs are predominantly located in intergenic regions (Fig. 1J) and that the chromosomal distribution of non-CG hyper DMRs were enriched in transposon-rich heterochromatins rather than in gene-rich euchromatic regions (Supplementary Figure 4). We examined methylation levels in both euchromatic and heterochromatic regions across fruit ripening stages (Supplementary Figure 13 and Supplementary Information). As expected, DNA methylation levels in fruit and leaf were higher in the heterochromatic regions than in the euchromatic regions across all three cytosine contexts. Notably, CHG and CHH methylation in heterochromatin is substantially higher in fruit compared to leaf throughout all five ripening stages—a pattern not observed in the euchromatic regions (Fig. 5A). These findings indicated that the regulation of CHG and CHH methylation in heterochromatin exhibits tissue-specific differences, with reproductive tissues displaying patterns different from those observed in somatic tissue.

**Fig. 5.**
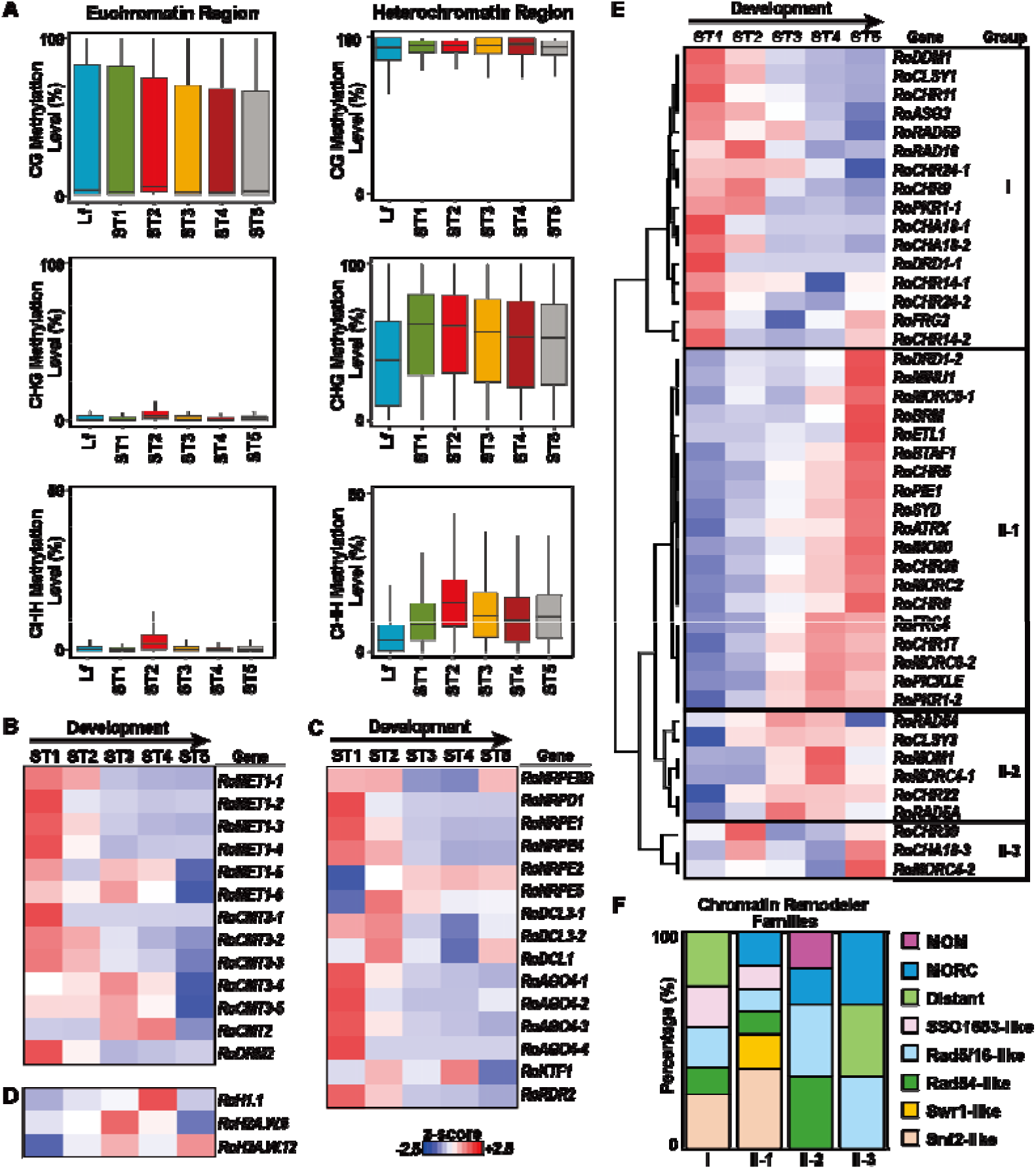
Euchromatic and heterochromatic DNA methylation and gene expression patterns of epigenetic-related genes during ripening. **A** DNA methylation levels in heterochromatic and euchromatic regions (50-bp windows) shown as box plots. Lf, leaf. Heatmaps present expression of genes related to (**B**) DNA methylation, (**C**) RNA-directed DNA methylation (RdDM) pathway, (**D**) histone variants, and (**E**) chromatin remodelers. **F** Chromatin remodeling genes are clustered into two major transcriptional groups across ripening stages by unsupervised hierarchical clustering analysis.

### Paradox of concurrent increase in non-CG methylation and transcripts of histone proteins associated with heterochromatin compaction

To understand the gene activities involved in catalyzing CHG and CHH methylation, we examined the transcript levels of DNA methyltransferase. We revealed that (i) CMT2, catalyzing CHH methylation in long transposons and heterochromatic transposons^27^, is active at the late fruit stages, suggesting that the increase in CHH methylation can, in part, be attributed to RoCMT2; (ii) RoDRM2 and most RdDM pathway genes, catalyzing CHH methylation in short transposons and euchromatic transposons^27^, are predominantly expressed during early stages; and (iii) RoCMT3, catalyzing CHG methylation, are active from early to late ripening stages (Fig. 5B, C and Supplementary Table 2C, 2D). These results suggested that a genome-wide increase in non-CG methylation within heterochromatin may be associated with elevated DNA methyltransferase transcript levels. However, heterochromatin is compact and less accessible. Therefore, increased heterochromatic DNA methylation during fruit ripening may reflect a shift toward greater chromatin accessibility in heterochromatin, a situation resembling that observed during *Arabidopsis* embryogenesis^28–30^. However, transcripts of H2A.W and H1—histone proteins known to promote chromatin compaction^31^—are increased during late ripening stages (Fig. 5D and Supplementary Table 2E). This paradox—elevated non-CG methylation alongside increased transcript levels contributing to heterochromatin compaction—may reflect a finely tuned regulatory epigenetic state whereby heterochromatin undergoes transient decondensation, allowing access by DNA methyltransferases despite the presence of compaction-associated histones.

### Multiple chromatin remodelers are active during black raspberry fruit ripening

To investigate the mechanisms underlying this enigma, i.e., elevated non-CG methylation with increased heterochromatin compaction, we analyzed the transcriptional profiles of chromatin remodelers that may facilitate heterochromatin opening. In *Arabidopsis*, the chromatin remodeler DDM1 is specifically required for cytosine methylation at H1-associated heterochromatin upon cell division^27^. However, cell division ceased after the early ripening stages and DDM1 was downregulated after ST2 (Fig. 5E and Supplementary Table 2F) in black raspberry, suggesting that additional chromatin remodelers might play roles in licensing the access of the methyltransferases to the heterochromatin to increase DNA methylation levels after ST2. To identify potential chromatin remodelers mediating heterochromatin accessibility after ST2, we conducted a homology-based gene search (Supplementary Information) and examined their expression patterns across fruit ripening. We identified 44 chromatin remodelers that are transcriptionally active during black raspberry fruit ripening and can be clustered into two major expression patterns (Fig. 5E and Supplementary Table 2F). Group I genes, including RoDDM1, were highly active during the early stages of fruit ripening, whereas group II genes were predominantly upregulated during the later stages. For example, a subgroup group II-1 consists of genes that were strongly expressed during the final stage ST5 (Fig. 5D). Interestingly, members of five gene families represented in group I also appear in group II (e.g., *RoCLSY* and *RoDRD* genes in RoRad54-like family), suggesting a possible subfunctionalization of family members resulting in a stage-preferred expression pattern during fruit ripening (Fig. 5D, F and Supplementary Table 2F). Furthermore, several gene families were uniquely represented in group II, including the *RoMOM* and *RoMORC* genes (Fig. 5F). Certain chromatin remodelers, such as MOM1, MORCs, CLSYs and DRDs^32,33^ are known to influence DNA methylation or direct RdDM in a locus-specific manner. Together, our results suggest a coordinated action among distinct chromatin remodelers that are active at different ripening stages to mediate chromatin accessibility to facilitate targeted DNA methylation reprogramming at specific genomic loci during black raspberry fruit ripening.

## Discussion

We observed that localized CG demethylation occurs primarily at loci in promoter regions where CHG and CHH are concurrently demethylated (Fig. 6). Promoter-associated methylation changes in genes, including TFs, are associated with changes in transcription and are enriched for functions pertinent to fruit ripening, such as key transcriptional regulators and hormone signaling pathways. DNA demethylases are known to act at specific loci to regulate gene activity. For example, the *Arabidopsis* DEMETER (DME) targets particular loci to induce hypomethylation and regulates imprinted gene expression in the endosperm^34^. However, the mechanisms by which DNA demethylases are recruited to these loci remain largely unknown. One of the major unresolved questions in fruit ripening is how DNA demethylases are targeted to specific loci within the fruit genome.

**Fig. 6.**
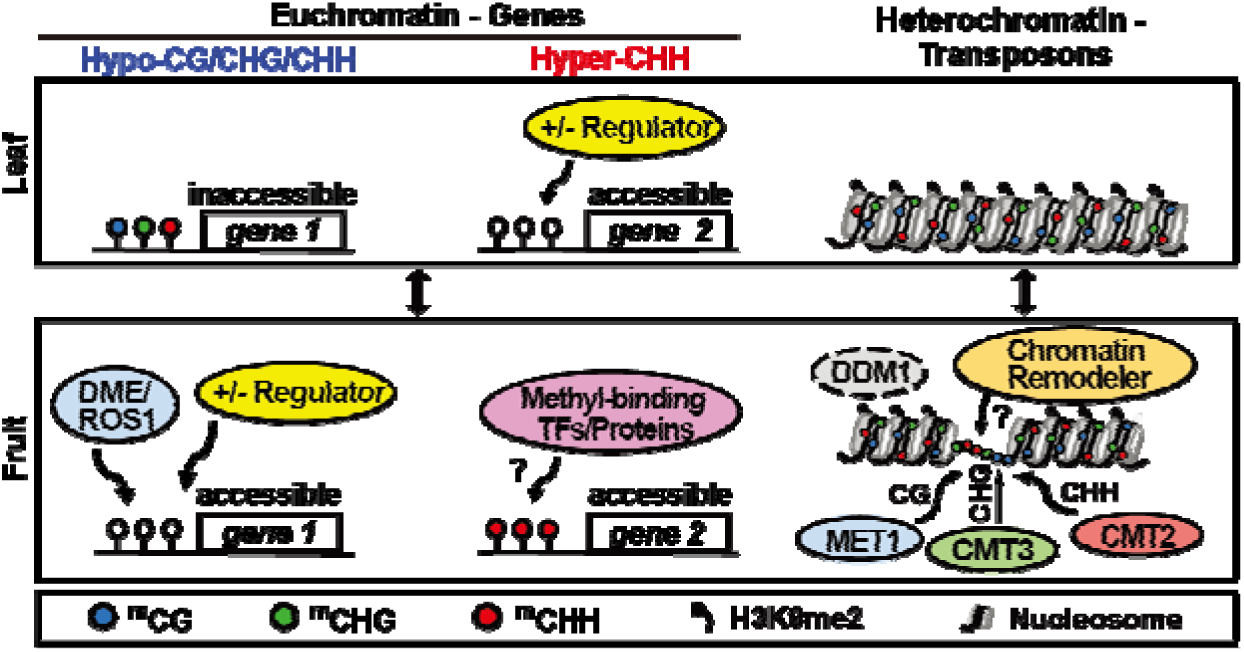
Schematic model of DNA methylation dynamics in euchromatic gene regions and heterochromatic transposons in the leaf and the fruit of black raspberry.

Our results demonstrate that CHH hypermethylation can be linked to both activation and repression of gene expression during black raspberry fruit ripening (Fig. 6), in agreement with strawberry methylome studies^14^. Factors involved in the transcription activation of methylated genes have been identified. For example, a 39 bp *cis*-element in the *Arabidopsis* DNA demethylase *ROS1* promoter functions as a methylation monitoring sequence (MEMS) and the DNA methylation level of the MEMS positively correlated with *ROS1* expression^35,36^. Subsequent studies on the MEMS and other methylated genes have identified a group of SUVH methyl-DNA binding proteins and their associated DNAJ domain transcriptional activators, which play roles in promoting the transcription of methylated genes^18,37^. In addition, during LEC2-induced somatic embryogenesis in *Arabidopsis*, the LEC2-activated RdDM pathway deposits CHH hypermethylation, which is recognized by the SUVH–SDJ reader complex that recruits AHL proteins to increase chromatin accessibility and activate totipotency-regulating genes^19^. By contrast, methylated DNA-mediated silencing is manifested through the recruitment of the *Arabidopsis* MBD5/6 complex that also contains the J-domain protein SILENZIO (SLN), and two α-crystalline domains (ACD) containing proteins ACD15 and ACD21 that promote higher-order multimerization of MBD5/6 complexes within heterochromatin ^38,39^. Furthermore, methyl-binding TFs such as the SHH homeodomain TF gene and the REM family genes (*VDD, VAL, REM12, REM13*), recruit RNA polymerase IV and direct DNA methylation at specific loci, particularly in reproductive tissues^40,41^. Together, these studies highlight how sequence-specific TFs control the spatial and temporal regulation of DNA methylation and provide a genetic framework for epigenetic patterning. One major challenge in understanding the role of CHH hypermethylation during fruit ripening is to identify what methyl-binding TFs or proteins are directed to methylated genes to drive specific transcriptional outputs. Our analysis of expression patterns of methyl-binding TFs and associated proteins may provide an important first step toward addressing this question.

Remarkably, relative to the leaf, black raspberry fruit methylomes show substantially elevated non-CG methylation. These changes, primarily confined to transposon-rich heterochromatin, indicate large-scale heterochromatin remodeling during fruit development. Divergent non-CG methylation patterns in reproductive and somatic tissues suggest tissue-specific regulatory mechanisms (Fig. 6), consistent with findings in endosperm, pollen and embryos^42,43^. One intriguing aspect of our results is the observation that many chromatin remodeler genes in fruit are dynamically regulated at transcriptional level at distinct stages, which may contribute to elevated non-CG methylation as compared to somatic tissues (Fig. 6), highlighting they are actively participating distinct biological functions in their respective stage of development. For example, *RoDDM1, RoCLSY1, RoDRD1* are highly active during early stages (ST1 and ST2; Fig 5E and Supplementary Table 2F) and contribute to the rapid increases in CHG and CHH methylation in these stages by facilitating chromatin accessibility to the RdDM machinery in the heterochromatin regions. By contrast, Group II chromatin remodelers (Fig 5E), active during ripening stages (ST4 & ST5), include RoBRM, RoPIE1, RoSYD, RoINO80, and RoPICKLE that may be involved in exchanging histone variants, altering nucleosome positioning, or epigenetically regulating gene transcription associated with the ripening stage of fruit development. Analysis of active chromatin remodeler genes during black raspberry fruit ripening reveals that, while some have been functionally characterized in prior studies, many others remain unexplored, leaving their contributions to ripening largely unknown. Clarifying their functions involved in this process remains an important outstanding question.

In summary, our results depict a dynamic, tightly regulated, and concerted molecular framework in which epigenomic modifications, transcriptional reprogramming, chromatin remodeling, and metabolic shifts act in coordination to orchestrate the black raspberry fruit ripening process.

## Methods

### Plant material

Black raspberry NC 349, an advanced selection from a Jewell x NC 100 cross developed by Dr. James Ballington, was planted at the North Carolina Department of Agriculture Piedmont Research Station in Salisbury, NC in 2012. Fruits and leaves used in this study were collected between June 10 and July 20, 2016. Black raspberry fruit development progresses through distinct color and firmness changes, from green to red, then violet, and finally black. Black raspberry leaves and fruits at five ripening stages were collected (Fig. 1) and bio-replicates of each were collected to make RNA-seq libraries, BS-seq libraries, and for the metabolome experiments.

### Reference genome used in this study

The reference genome of *Rubus occidentalis* v3.0 was downloaded from Genome Database for Rosaceae (GDR, https://www.rosaceae.org/analysis/268), for data processing and analysis.

### BS-Seq Library Construction, Methylome Sequencing, Data Processing, and Sequence Analysis

Fruits of five ripening stages (Fig. 1) with two bio-replicates were collected to extract genomic DNA as subject for sodium bisulfite treatment followed by BS-seq library construction. In addition, genomic DNA extracted from leaves was also used for BS-seq library construction as a control to compare methylation levels with fruits. These BS-seq libraries were sequenced using Illumina platform with 150-base paired-end reads. DNA was prepared for BS-Seq library preparation and methylome sequencing following the methods of previous published methods^44^. Details are provided in the Supplementary Information.

### DMR analysis by CGmapTools

To identify the changes in local DNA methylation levels, which is called differentially methylated regions (DMR) during black raspberry fruit ripening. We applied CGmapTools^16^ to identify DMRs with the cutoff *p* < 0.05, CG > 20%, CHG > 10% and CHH > 8%. Moreover, the identified two or more DMRs whose distance are less than 300 bps are merged into one merged-DMR. To precisely and comprehensively inspect the occurrence of DMRs, DNA methylation levels in leaves were used as baseline reference for comparison with those in each fruit ripening stage.

### RNA-Seq Library Construction, Sequencing, Data Processing, and Analysis

Three bio-replicates each of black raspberry fruits were collected at the five ripening stages. Total RNA was isolated from fruits and leaves, using the RNeasy Plant Mini Kit (Qiagen) and treated with RNase-free DNase I (Qiagen). Total RNA with a minimum RIN (RNA Integrity Number) value of 8 was employed for RNA-Seq library construction. Poly-A+ RNA was selected using the NEBNext Poly(A) mRNA Magnetic Isolation Module (NEB) for RNA-Seq library construction using the KAPA Stranded mRNA-Seq Kit (Roche). Previously published methods were applied for data processing and downstream analysis, including WGCNA analysis^45^. Details are provided in the Supplementary Information.

### Metabolome profiling

Three biological replicates of fruits at the five ripening stages (Fig. 1) were collected to extract metabolites for high-performance liquid chromatography with diode-array detection (HPLC-DAD), following previously published methods^46^. Briefly, filtered samples were injected (10 µL) into a 1200 HPLC system (Agilent Technologies, Santa Clara, CA, USA) equipped with a UV–vis diode array detector (DAD), controlled-temperature autosampler (4°C), and column compartment (30°C) using a reversed-phase Supelcosil LC-18 column, 25 mm × 4.6 mm × 5 µm (Supelco, Bellefonte, PA, USA). Standard curves were calculated using peak areas at UV of 520 nm as cyaniding-3-O-glucoside equivalents.

### Statistics and reproducibility

For each cytosine in the reference genome, we quantified the number of reads reporting cytosines (methylated cytosines) and thymines (resulting from bisulfite conversion of unmethylated cytosines), using custom scripts. Sites with only a single mapped read (either cytosine or thymine) were excluded to reduce noise. To further minimize false positives, we applied a binomial test (*p* < 0.05) to determine whether the observed methylation at a site was statistically significant. Cytosines passing this threshold—typically those with at least two cytosine reads—were considered methylated. Conversely, cytosine sites with at least one thymine read and failing the binomial test were considered unmethylated, indicating that bisulfite conversion of unmethylated cytosines to uracils (detected as thymines in sequencing). Average methylation level was defined as the percentage of methylated cytosines across all filtered sites and calculated using the formula: [C / (C + T)] × 100, where C represents the number of methylated reads and T represents unmethylated reads. This calculation was performed separately for cytosines in CG, CHG, and CHH sequence contexts. To estimate methylation levels over specific genomic features, we computed the mean of methylation levels for all cytosine sites within the feature.

To explore the global DNA methylation differences of leaf stage and five fruit ripening stages the Student’s t-test was used. The associations between gene modules and phenotypic traits (metabolite data), Pearson’s correlation was calculated between module eigengenes and sample traits, and the significance of each correlation was assessed by Student’s t-test.

For the reproducibility of 12 biological replicates of BS-Seq data across leaf stage and five fruit ripening stages, the correlation coefficient was applied to the average methylation levels in 500-kb windows across the genome from biological replicates. For the reproducibility of 14 biological replicates of RNA-Seq data, principal component analysis (PCA) was performed on transcript data from five fruit ripening stages.

## Acknowledgements

This work is supported by the National Institute of Food and Agriculture Hatch Projects NC02413 and NC02891 (to T.-F. H.), NC02895 (to X. L.), Southern Small Fruits Consortium Grant 2010-01 (to P.P.-V), an internal PHHI Seed Grant to (T.-F. H., X. L., and P. P.-V.), the Agricultural Biotechnology Research Center of Academia Sinica (to J.-Y. L.), and the Biotechnology Research Center in Southern Taiwan of Academia Sinica (to J.-Y. L.).

## Contributions

T.-F. H., X. L. and J.-Y. L. designed the research; Y.-H. H., J.-H. Z., H.-Y. C. and P. P.-V. performed vthe experiments; G. F. provided research materials; W.-H. H., M.-H. H., L.-P. L., Y.-C. W., Y.-H. H., B.H. L., Y.-Y. H., J.-H. Z and H.-Y. C. analyzed the data; J.-Y. L., T.-F. H. and W.-H. H. revised the manuscript.

## Data availability statement

The raw and processed RNA-seq data reported in this paper has been uploaded to NCBI (accession number: GSE).

