## Supplementary Information for "Multi-Omics analyses reveal dynamic interactions between DNA methylation and transcriptional regulation during black raspberry ripening"

### Supplementary Methods

#### BS-Seq library construction, methylome sequencing, data processing, and sequence analysis

DNA sequences were aligned to the black raspberry whole genome assembly and annotation (version 3.0; https://www.rosaceae.org/analysis/268). To assess bisulfite (BS) conversion efficiency, 3 ng of unmethylated cI857 Sam7 λ DNA (GenBank Accession No. NC_001416; Promega) was spiked into the genomic DNA prior to sonication as an internal control. Adapter-ligated DNA libraries underwent two rounds of BS treatment using the EZ DNA Methylation-Lightning Kit (Zymo Research). Following BS conversion, DNA was purified and amplified for 10 cycles using Q5U DNA polymerase (NEB). Size selection of PCR-amplified fragments was performed using AMPure XP beads (Beckman). Sequencing reads were aligned to the reference genome using BS-Seeker2[1], allowing up to four mismatches. Cytosine coverage was calculated as the total number of cytosines (methylated) and thymines (converted/unmethylated) detected across all reads. Only uniquely mapped reads were retained for downstream analysis. Postprocessing of the BS-Seeker2 output involved two quality control steps: (i) removal of PCR duplicates by collapsing reads with identical 5′-end mapping positions and nucleotide sequences, and (ii) filtering out reads containing three or more consecutive cytosines in the CHH context, which are indicative of incomplete BS conversion[2].

#### Assessment of bisulfite conversion efficiency

To evaluate the efficiency of bisulfite (BS) conversion and estimate the non-conversion rate, we included unmethylated λ DNA as an internal control by spiking it into each genomic DNA sample for library preparation. Sequencing yielded a minimum of 300× coverage across the λ genome, allowing for robust analysis. We detected more than 99% of C-to-T conversion within the λ genome, indicating highly efficient BS conversion.

#### Identification of differentially methylated genes by colocalization analysis

We defined differentially methylated genes (DMGs) as genes that have differentially methylated regions (DMRs) in the promoter (1.5 Kb upstream of the transcription start site) and gene body. We scanned the genome using the intersect function of BEDTools[3] with coordinates from DMRs and all genes in the black raspberry genome to colocalize DMRs and genes.

**Identification of genes whose transcriptional activity correlates with alterations in DNA methylation**

DMGs and differentially expressed genes (DEGs) were integrated to investigate the relationship between DNA methylation and gene expression. Genes that were both differentially methylated and differentially expressed were identified as genes whose transcriptional activity is regulated by DNA methylation.

**RNA-Seq library construction, sequencing, data processing, and analysis**

RNA-seq libraries were sequenced using the Illumina platform to generate 150-base paired-end reads. Illumina raw reads were processed with Trimmomatic[4] to eliminate adapters and to trim low-quality bases at the 5' and 3' ends, targeting positions with error rates greater than 0.5% and error rate > 0.1%. Previously published methods were applied to process RNA-Seq data with modifications[2]. The remaining high-quality reads were mapped to the reference genome using Hisat2[5] with mismatch 4. Only uniquely mapped reads (i.e., reads that map to one unique genomic locus) were used for subsequent analysis. The read counts per gene were analyzed using HTSeq[6], followed by using DESeq2[7] to determine DEGs with cutoff $q$-value < 0.05 and > two-fold changes. Non-supervised hierarchical clustering was analyzed by dChip[8]. To generate a comprehensive Gene Ontology (GO) database for black raspberry, we used predicted protein sequences and Blast2GO[9] with BLASTP for homologous research. This approach enabled the assignment of functional GO terms to the predicted protein-coding genes to construct a GO database. The goseq package[10] was used for gene GO enrichment analysis with a cutoff FDR (Benjamini–Hochberg multiple testing correction) < 0.05.

**Weighted gene correlation networks for analysis (WGCNA) analysis**

The WGCNA R package[11] was employed to identify gene coexpression modules of highly correlated expressed genes using DESeq2 normalized data by following published methods[12] with modifications. We first plotted the distribution of RNA abundance for each ripening stage using a density plot based on $\log_2$(DESeq2 normalized count + 1) values. We identified the lowest point between the two peaks of the bimodal distribution, which represents the threshold for low expression. Genes with expression levels below this point were considered to be expressed at low levels and excluded from the WGCNA analysis. The lowest points for the five ripening stages (from green to black) were 3.06, 3.49, 3.12, 3.16, and 3.91, respectively. Finally, the lowest value (3.06) among the five stages was used as the cutoff across all stages. In the initial step of WGCNA, an appropriate soft-thresholding power is selected to define the network's scale-free topology. This selection is guided by the $R^2$ criterion, which helps identify the power value that best balances network connectivity and biological relevance in the weighted adjacency matrix. The adjacency matrix was converted into a topological overlap matrix (TOM) to evaluate the similarity of gene connectivity. The TOM-based dissimilarity was then used for module identification through hierarchical clustering, and module eigengenes—defined as the first principal component of each module—were computed. Modules with highly similar expression profiles were merged. To explore the associations between gene modules and phenotypic traits (metabolite data), Pearson's correlation was calculated between module eigengenes and sample traits, and the significance of each correlation was assessed using Student's t-test.

**Homology search to deduce enzyme-coding genes in anthocyanin biosynthesis**

With respect to the metabolite profiles and the addition of functional groups to the substrates, we predicted enzymes that catalyze the transfer reaction to produce the seven anthocyanins (Fig. 2), including anthocyanidin 3-O-glucoside 2-O-xylosyltransferase, anthocyanin 3-O-glucoside rhamnosyltransferase, anthocyanidin 3-O-glucoside xylosyltransferase, anthocyanidin 3-O-glucosyltransferase and anthocyanin O-methyltransferase. We conducted a protein homology search using protein sequences from other plant species with experimentally proven enzyme functions to BLASTP against the black raspberry protein database to identify candidate genes with the most significant E-values in the black raspberry genome.

**Identification of anthocyanin- and ripening-related transcription factors in black raspberry**

To identify putative transcription factors (TFs) involved in fruit ripening and anthocyanin biosynthesis in black raspberry, we first searched publications and review articles related to these processes across various plant species (Supplementary Table 7). According to these previous studies, we selected TFs that were either functionally validated or widely recognized in the literature. Subsequently, the protein sequences of these known TFs were retrieved and subjected to BLASTP analysis against the protein database of black raspberry. For each query sequence, the hit with the lowest E-value (threshold: $1 \times 10^{-4}$) was selected as the best match. In cases where multiple query TFs aligned to the same black raspberry TF, we retained the one with the most significant E-value. If two or more hits had identical E-values, the TF from the species phylogenetically closest to black raspberry or with the highest protein sequence identity was selected as the representative. This approach ensured that each well-characterized TF from other species was systematically assigned to a corresponding blackberry homolog with the highest confidence.

**Sampling heterochromatic and euchromatic regions of black raspberry genome**

The euchromatin and heterochromatin regions were randomly selected through DNA methylation profiling, transposon and gene density across seven chromosomes of black raspberry by Integrative Genomics Viewer v.2.19.4.[13]. The region with a high level of DNA methylation, high transposon density and low gene density was defined as the heterochromatin region. Conversely, the region with a low level of DNA methylation, low transposon density and high gene density was defined as the euchromatin region.

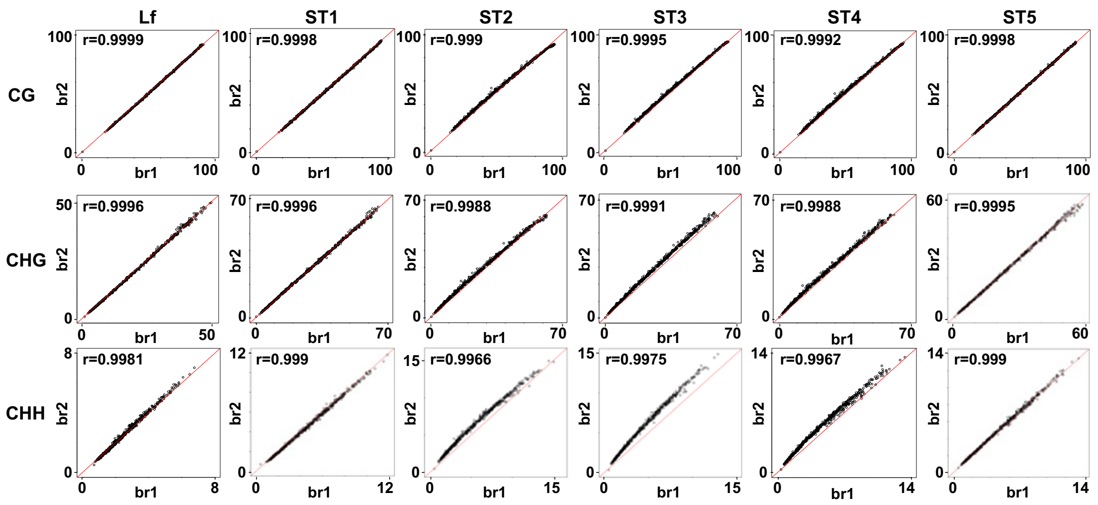


**Supplementary Figure 1 | Quality of BS-Seq methylome libraries. Correlation coefficients between biological replicates of black raspberry BS-Seq libraries.** The average methylation levels in 500-kb windows across the genome from biological replicates with similar sequencing depths were used to determine the correlation coefficients. Lf, leaf; ST, stage.

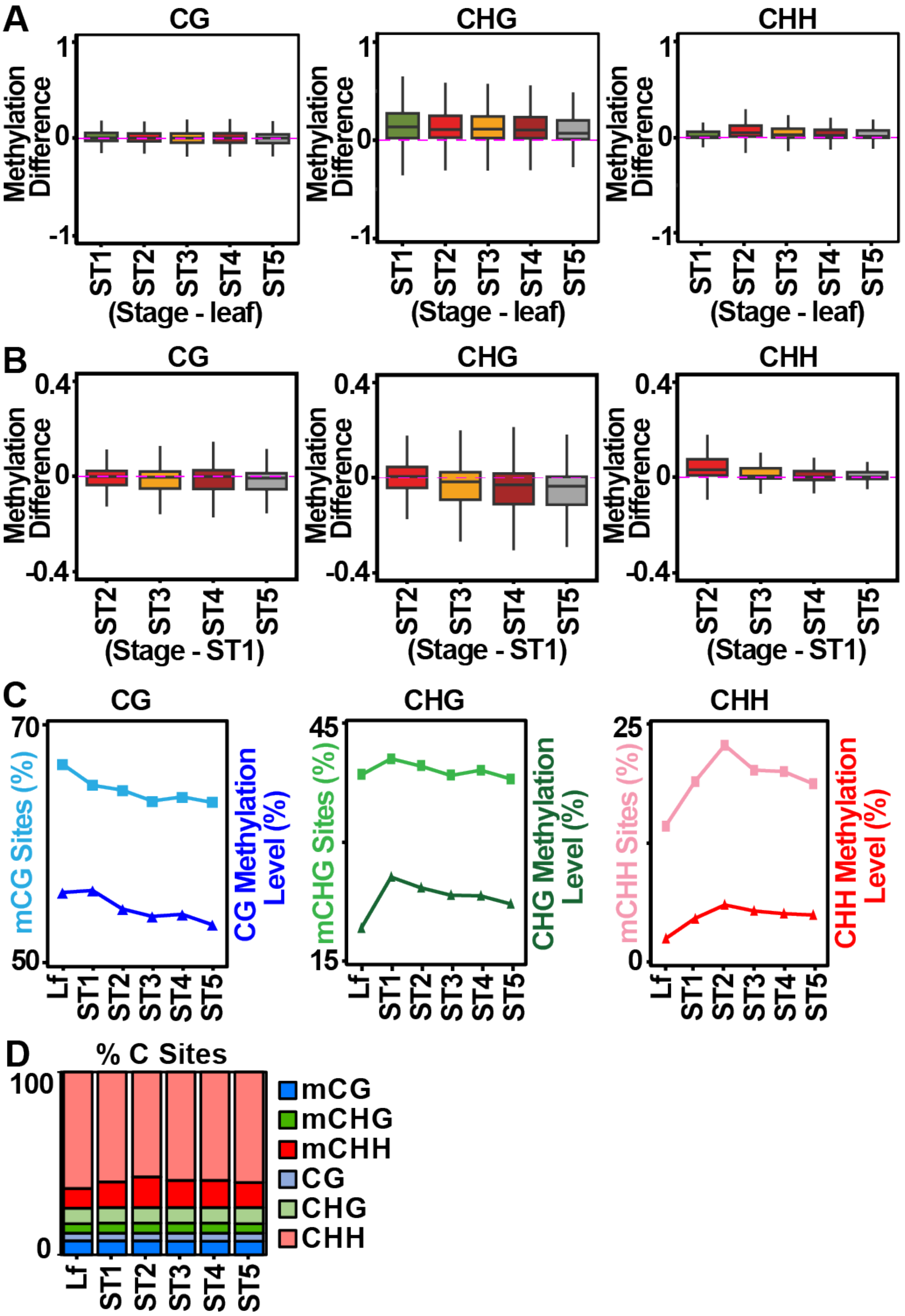
A
CG
CHG
CHH
Methylation Difference
1
0
-1
ST1
ST2
ST3
ST4
ST5
(Stage - leaf)
B
Methylation Difference
0.4
0
-0.4
(Stage - ST1)
C
mCG Sites (%)
CG Methylation Level (%)
mCHG Sites (%)
CHG Methylation Level (%)
mCHH Sites (%)
CHH Methylation Level (%)
70
50
45
15
25
0
Lf
D
% C Sites
100
mCG
mCHG
mCHH
CG
CHG
CHH

**Supplementary Figure 2 | DNA methylation dynamics during black raspberry fruit ripening. A** Genome-wide DNA methylation levels in fruit versus leaf tissues analyzed in 50-bp windows. **B** Differential DNA methylation between late ripening stages versus stage 1 calculated using 50-bp windows. **C** DNA methylation changes during black raspberry fruit development assessed by comparing the proportion of methylated cytosine sites and the average methylation percentages. The latter represents the overall mean DNA methylation levels across all cytosine sites in the black raspberry genome. **D** Proportions of cytosine bases in CG, CHG, and CHH contexts for the black raspberry fruit genome. Lf, leaf; ST, stage.

A

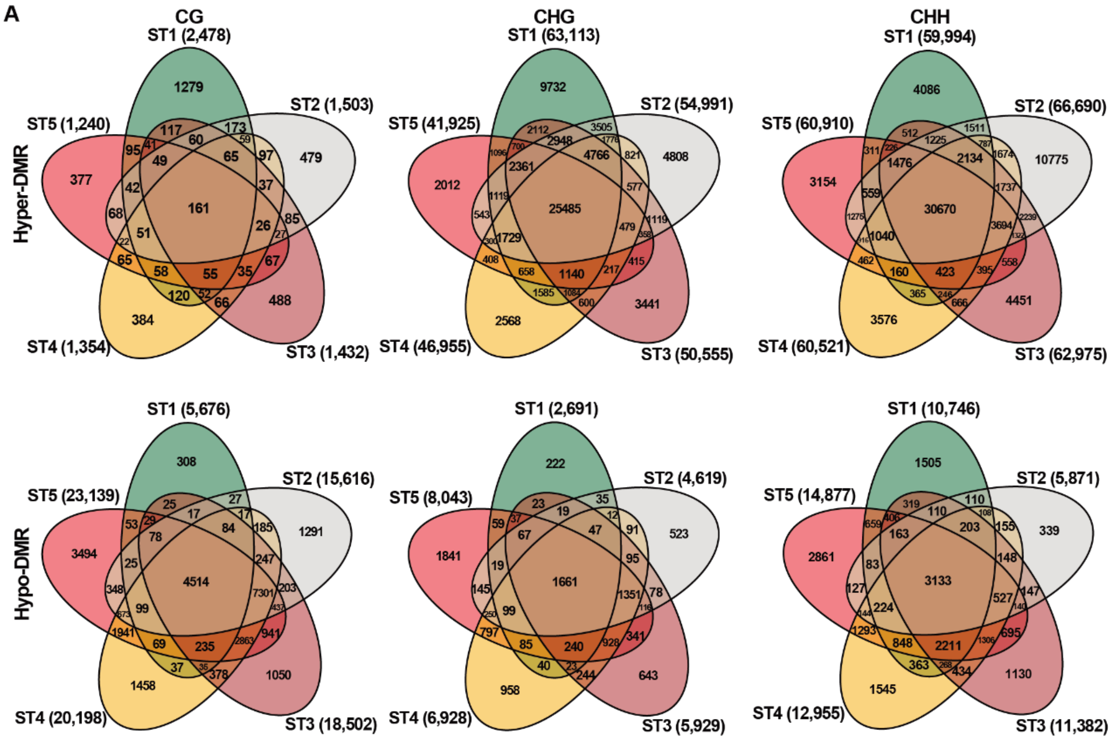


B

| | CG-hyper-DMR | | | | CHG-hyper-DMR | | | | CHH-hyper-DMR | | | |
|---|---|---|---|---|---|---|---|---|---|---|---|---|
| Stage | Non-unique hyper DMR | Unique hyper DMR | Percentage of overlap | Total | Non-unique hyper DMR | Unique hyper DMR | Percentage of overlap | Total | Non-unique hyper DMR | Unique hyper DMR | Percentage of overlap | Total |
| ST1 | 1,199 | 1,279 (51.6%) | 6.5% | 2,478 | 53,381 | 9,732 (15.4%) | 40.6% | 63,113 | 55,908 | 4,086 (6.8%) | 53.3% | 59,994 |
| ST2 | 1,024 | 479 (31.9%) | 10.7% | 1,503 | 50,183 | 4,808 (8.7%) | 46.6% | 54,991 | 55,915 | 10,775 (16.2%) | 47.9% | 66,690 |
| ST3 | 944 | 488 (34.1%) | 11.2% | 1,432 | 47,114 | 3,441 (6.8%) | 50.6% | 50,555 | 58,524 | 4,451 (7.1%) | 50.7% | 62,975 |
| ST4 | 970 | 384 (28.4%) | 11.9% | 1,354 | 44,387 | 2,568 (5.5%) | 54.5% | 46,955 | 56,945 | 3,576 (5.9%) | 52.8% | 60,521 |
| ST5 | 863 | 377 (30.4%) | 13.0% | 1,240 | 39,913 | 2,012 (4.8%) | 61.1% | 41,925 | 57,756 | 3,154 (5.2%) | 52.5% | 60,910 |
| Overlap | 161 | | | | 25,603 | | | | 31,951 | | | |
| | CG-hypo-DMR | | | | CHG-hypo-DMR | | | | CHH-hypo-DMR | | | |
| Stage | Non-unique hypo DMR | Unique hypo DMR | Percentage of overlap | Total | Non-unique hypo DMR | Unique hypo DMR | Percentage of overlap | Total | Non-unique hypo DMR | Unique hypo DMR | Percentage of overlap | Total |
| ST1 | 5,368 | 308 (5.4%) | 79.5% | 5,676 | 2,469 | 222 (8.2%) | 61.7% | 2,691 | 9,241 | 1,505 (14.0%) | 29.2% | 10,746 |
| ST2 | 14,325 | 1,291 (8.3%) | 28.9% | 15,616 | 4,096 | 523 (11.3%) | 36.0% | 4,619 | 5,532 | 339 (5.8%) | 53.4% | 5,871 |
| ST3 | 17,452 | 1,050 (5.7%) | 24.4% | 18,502 | 5,286 | 643 (10.8%) | 28.0% | 5,929 | 10,252 | 1,130 (9.9%) | 27.5% | 11,382 |
| ST4 | 18,740 | 1,458 (7.2%) | 22.3% | 20,198 | 5,970 | 958 (13.8%) | 24.0% | 6,928 | 11,410 | 1,545 (11.9%) | 24.2% | 12,955 |
| ST5 | 19,645 | 3,494 (15.1%) | 19.5% | 23,139 | 6,202 | 1,841 (22.9%) | 20.7% | 8,043 | 12,016 | 2,861 (19.2%) | 21.1% | 14,877 |
| Overlap | 4,514 | | | | 1,661 | | | | 3,133 | | | |

**Supplementary Figure 3 | Overlap of differentially methylated regions (DMRs) among black raspberry fruit ripening stages using leaf tissues as the reference. A** The number of DMRs shown as Venn diagrams. **B** The number and percentage of DMRs that are unique to a single stage or shared across more than one stage (non-unique).

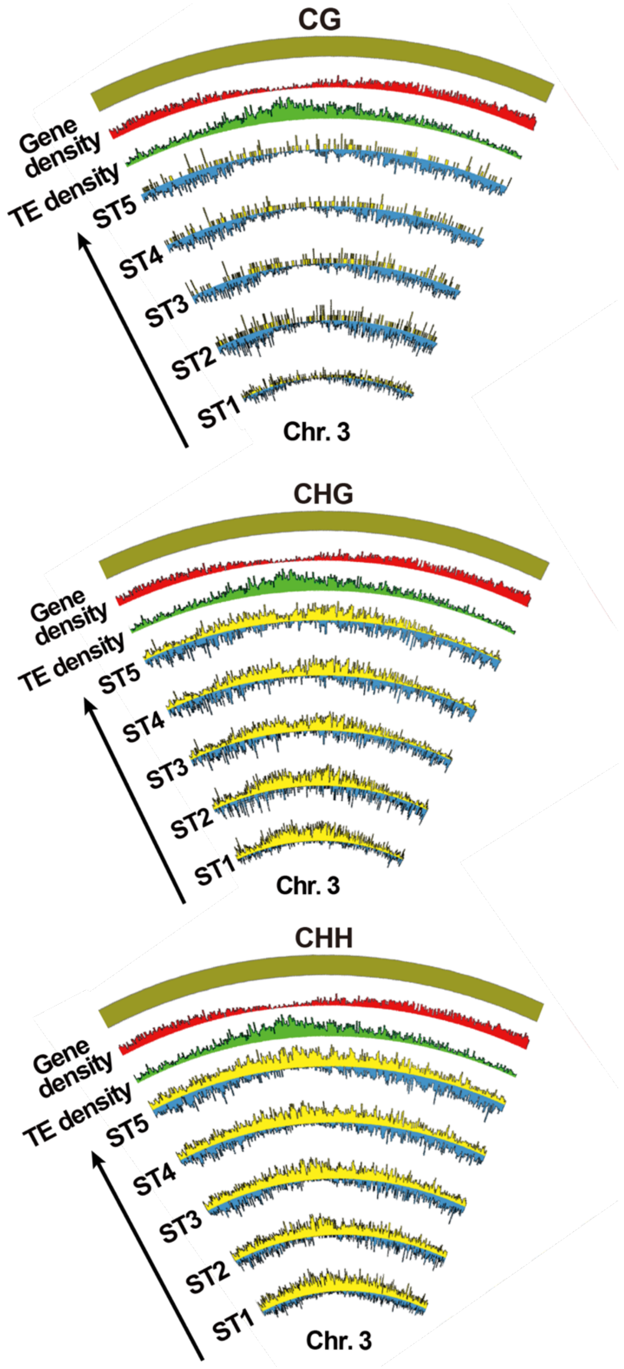


**Supplementary Figure 4 | Local DNA methylation changes during black raspberry fruit ripening.** The distribution of differentially methylated regions (DMRs) across chromosome 3, using leaf as the reference, is shown as an example. Hypo-methylated DMRs are shown in blue, and hyper-methylated DMRs are shown in yellow. Gene (red) and transposable element (TE; green) tracks represent the densities of genes and LTR retrotransposons along the chromosome.

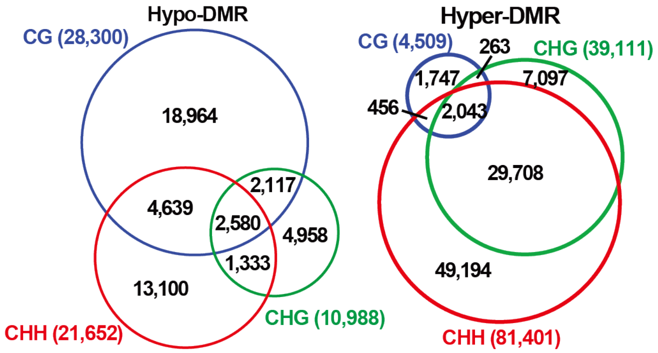


**Supplementary Figure 5 | Overlap of differentially methylated regions (DMRs) across CG, CHG, and CHH contexts using leaf as reference.**

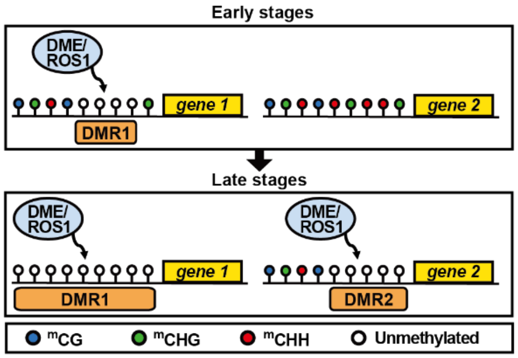


**Supplementary Figure 6 | Illustration of the spatial and temporal progression of DNA demethylation during fruit ripening.** Specific loci are demethylated at the early stages and gradually expand, while others maintain methylation until later stages.

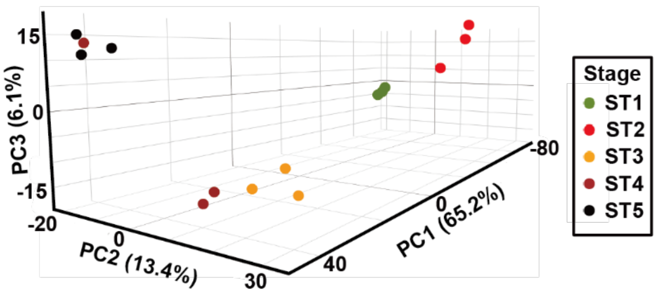


**Supplementary Figure 7 | Genome-wide gene expression patterns during black raspberry fruit ripening.** Principal Component Analysis (PCA) was performed on transcript data from different fruit stages. Principal Components 1 through 3 (PC1–PC3) together accounted for 85% of the variance in mRNA expression across the five stages.

| Category | GO terms | Green->Red | | Red->Soft | | Soft->Turning | | Turning->Black | |
|---|---|---|---|---|---|---|---|---|---|
| | | up | down | up | down | up | down | up | down |
| Flavonoid metabolic process | flavonoid biosynthetic process | | | | | | | | |
| Carbohydrate Related | glycolytic process | | | | | | | | |
| | carbohydrate metabolic process | | | | | | | | |
| Catalytic Activity | negative regulation of catalytic activity | | | | | | | | |
| Lipid Related | lipid metabolic process | | | | | | | | |
| | sterol metabolic process | | | | | | | | |
| | fatty acid biosynthetic process | | | | | | | | |
| Cell Cycle | regulation of cyclin-dependent protein serine/threonine kinase activity | | | | | | | | |
| | mitotic cell cycle phase transition | | | | | | | | |
| | DNA duplex unwinding | | | | | | | | |
| | cell division | | | | | | | | |
| | mitotic chromosome condensation | | | | | | | | |
| | DNA replication checkpoint signaling | | | | | | | | |
| Cell Wall Related | cellulose biosynthetic process | | | | | | | | |
| | lignin catabolic process | | | | | | | | |
| | plant-type secondary cell wall biogenesis | | | | | | | | |
| | arabinan catabolic process | | | | | | | | |
| Cytoskeleton | microtubule-based movement | | | | | | | | |
| Development | developmental process | | | | | | | | |
| | abaxial cell fate specification | | | | | | | | |
| | microtubule-based process | | | | | | | | |
| Energy Related | coenzyme A metabolic process | | | | | | | | |
| | mitochondrion organization | | | | | | | | |
| Photosynthesis | chlorophyll biosynthetic process | | | | | | | | |
| | photosynthetic electron transport in photosystem I | | | | | | | | |
| | NADH dehydrogenase complex (plastoquinone) assembly | | | | | | | | |
| | photosynthesis | | | | | | | | |
| | photorespiration | | | | | | | | |
| | formaldehyde catabolic process | | | | | | | | |
| Epigenetics | methylation | | | | | | | | |
| | chromatin silencing | | | | | | | | |
| | histone H3-K4 demethylation, trimethyl-H3-K4-specific | | | | | | | | |
| | histone H3-K36 demethylation | | | | | | | | |
| | C-5 methylation of cytosine | | | | | | | | |
| Hormone Related | brassinosteroid homeostasis | | | | | | | | |
| | auxin-activated signaling pathway | | | | | | | | |
| | abscisic acid-activated signaling pathway | | | | | | | | |
| Protein Modification Process | protein complex oligomerization | | | | | | | | |
| | protein-chromophore linkage | | | | | | | | |
| | dephosphorylation | | | | | | | | |
| | protein phosphorylation | | | | | | | | |
| Response To Stimulus | response to hydrogen peroxide | | | | | | | | |
| | response to heat | | | | | | | | |
| | response to oxidative stress | | | | | | | | |
| | response to light stimulus | | | | | | | | |
| | trehalose biosynthetic process | | | | | | | | |
| | xenobiotic detoxification by transmembrane export across the plasma membrane | | | | | | | | |
| Salvage & Protection | protein folding | | | | | | | | |
| | hydrogen peroxide catabolic process | | | | | | | | |
| | cellular oxidant detoxification | | | | | | | | |
| DNA Repair | DNA repair | | | | | | | | |
| | double-strand break repair via homologous recombination | | | | | | | | |
| | double-strand break repair via break-induced replication | | | | | | | | |
| | interstrand cross-link repair | | | | | | | | |
| Transport | zinc ion transmembrane transport | | | | | | | | |
| | amino acid transport | | | | | | | | |
| | water transport | | | | | | | | |
| | transmembrane transport | | | | | | | | |
| | dipeptide transmembrane transport | | | | | | | | |
| RNA Modification | RNA modification | | | | | | | | |
| | cytidine to uridine editing | | | | | | | | |
| | chloroplast RNA modification | | | | | | | | |
| | mitochondrial mRNA modification | | | | | | | | |
| RNA Processing | rRNA processing | | | | | | | | |
| DNA Metabolic Process | DNA biosynthetic process | | | | | | | | |
| Transcription | positive regulation of transcription, DNA-templated | | | | | | | | |
| | regulation of transcription, DNA-templated | | | | | | | | |
| | DNA-templated transcription, termination | | | | | | | | |
| Translation | plastid translation | | | | | | | | |

1.3 $-\log_{10}$(FDR) 30

**Supplementary Figure 8 | Gene Ontology (GO) analysis of differentially expressed genes (DEGs) between consecutive stages of black raspberry fruit ripening.** Representative enriched GO terms are shown, and the full list is available in Supplementary Table 3. “Up” and “Down” indicate genes that are upregulated or downregulated, respectively, between adjacent ripening stages.

| | CG-hyper-DMR | | CHG-hyper-DMR | | CHH-hyper-DMR | |
|---|---|---|---|---|---|---|
| ST1 base | RNA down | RNA up | RNA down | RNA up | RNA down | RNA up |
| ST2 | 29 | 31 | 84 | 45 | 1,175 | 593 |
| ST3 | 58 | 31 | 100 | 62 | 1,184 | 679 |
| ST4 | 52 | 35 | 76 | 51 | 805 | 482 |
| ST5 | 55 | 30 | 64 | 38 | 874 | 528 |
| | CG-hypo-DMR | | CHG-hypo-DMR | | CHH-hypo-DMR | |
| ST1 base | RNA down | RNA up | RNA down | RNA up | RNA down | RNA up |
| ST2 | 786 | 526 | 410 | 263 | 372 | 197 |
| ST3 | 1,851 | 1,291 | 990 | 578 | 1,222 | 747 |
| ST4 | 2,159 | 1,602 | 1,144 | 769 | 1,540 | 1,016 |
| ST5 | 2,820 | 1,956 | 1,537 | 1,007 | 2,082 | 1,375 |

**Supplementary Figure 9 | Number of genes identified as both differentially expressed (DEGs) and differentially methylated (DMGs).** "RNA down" and "RNA up" indicate genes that are downregulated or upregulated, respectively, relative to the green stage (ST1).

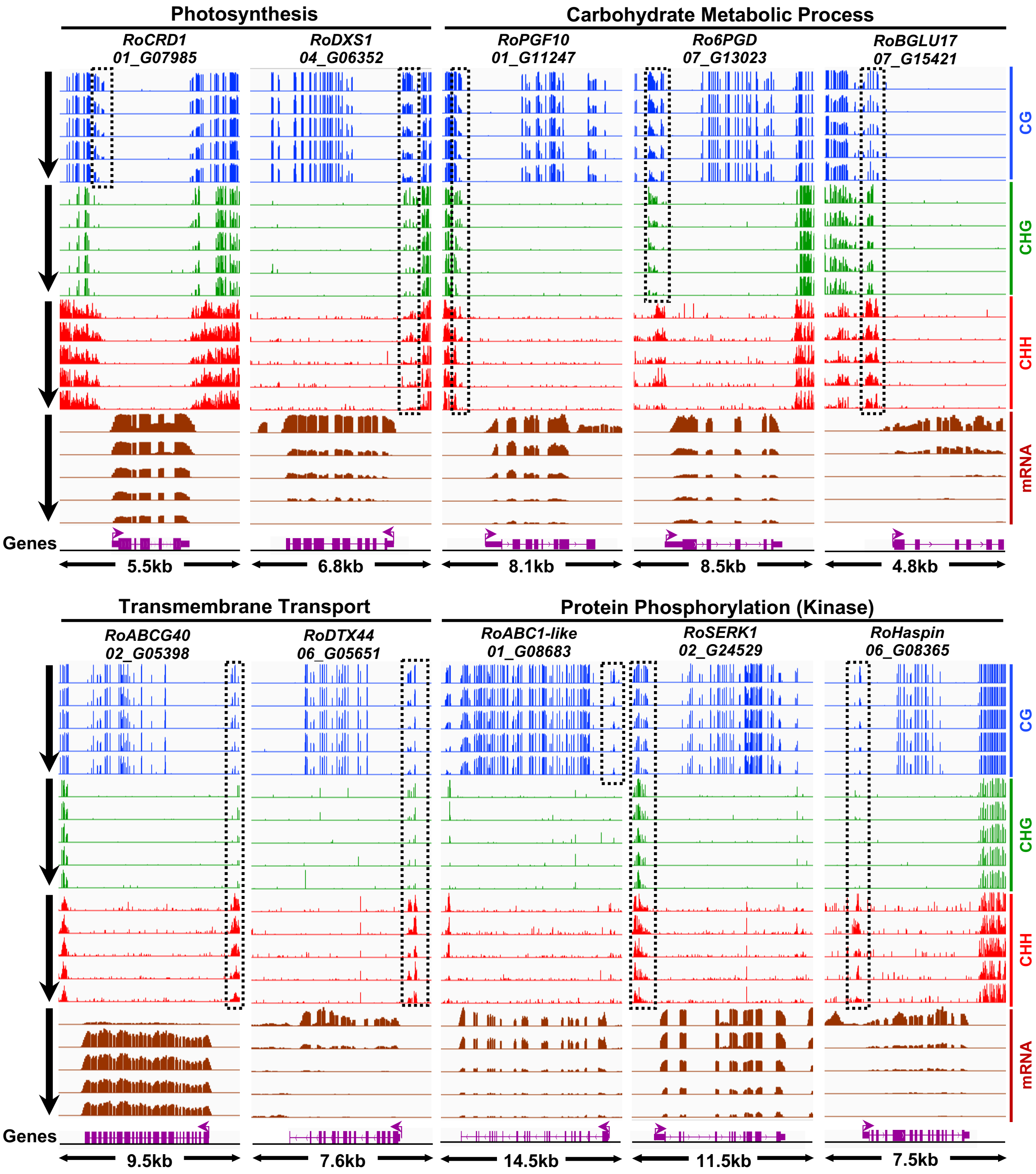

Photosynthesis
Carbohydrate Metabolic Process
RoCRD1
01_G07985
RoDXS1
04_G06352
RoPGF10
01_G11247
Ro6PGD
07_G13023
RoBGLU17
07_G15421
CG
CHG
CHH
mRNA
Genes
5.5kb
6.8kb
8.1kb
8.5kb
4.8kb
Transmembrane Transport
Protein Phosphorylation (Kinase)
RoABCG40
02_G05398
RoDTX44
06_G05651
RoABC1-like
01_G08683
RoSERK1
02_G24529
RoHaspin
06_G08365
9.5kb
7.6kb
14.5kb
11.5kb
7.5kb

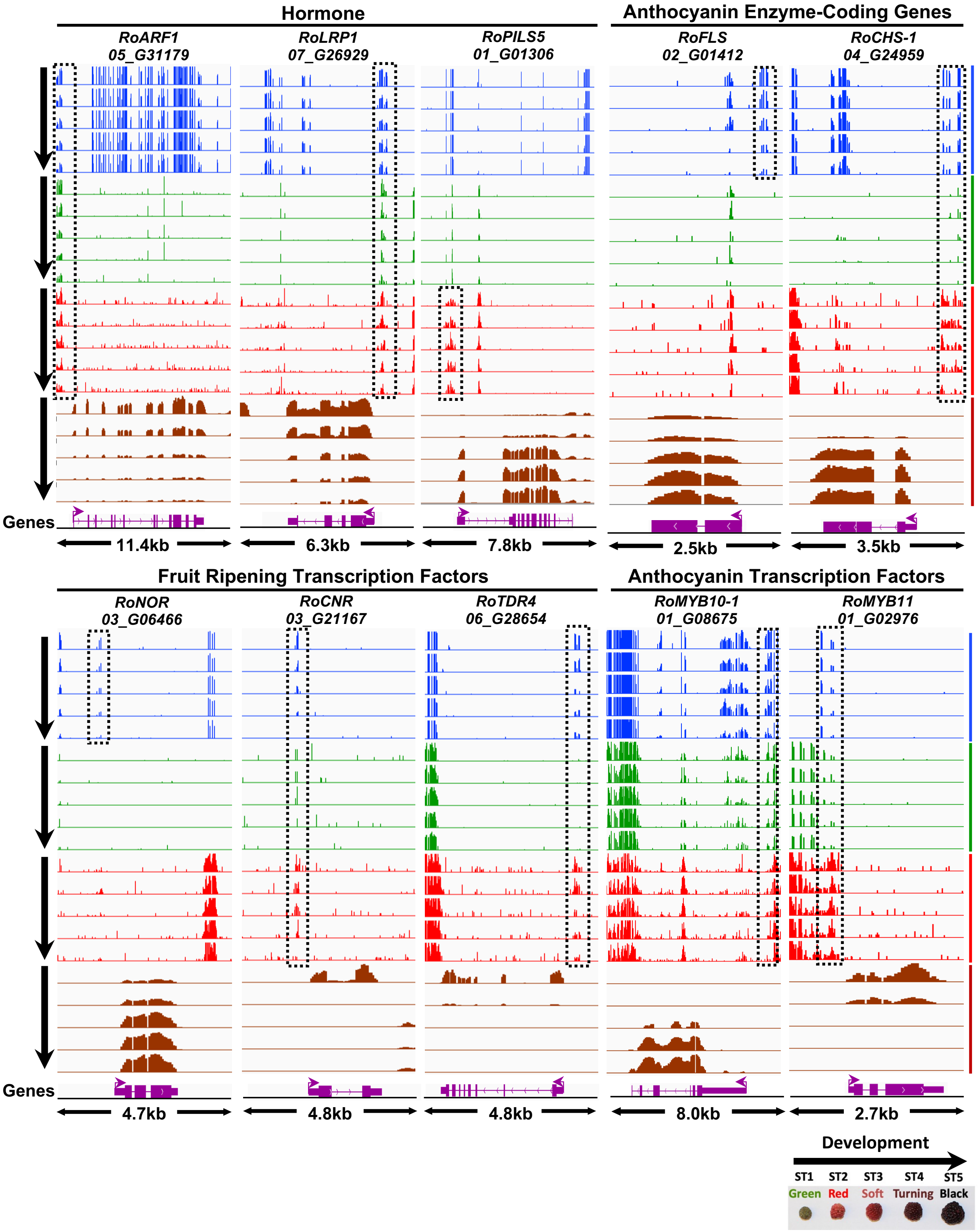


**Supplementary Figure 10 | Methylation levels and mRNA accumulation patterns of gene classes during black raspberry fruit ripening.**

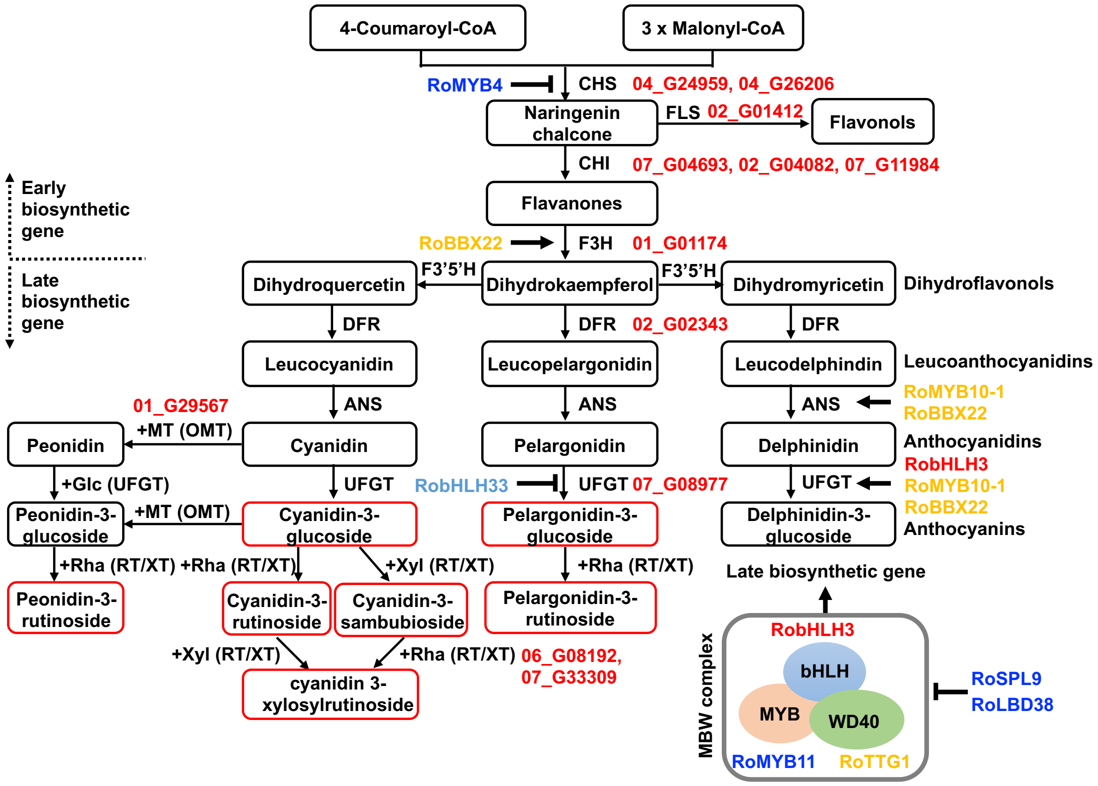


**Supplementary Figure 11 | Diagram of the anthocyanin biosynthetic pathway in black raspberry fruit.** Gene symbols are color-coded based on the Pearson correlation between transcript abundance and relative anthocyanin content across developmental stages. Red indicates strong positive correlation ($r \geq 0.8$), and blue indicates strong negative correlation ($r \leq -0.8$). Orange and light blue indicate moderate positive ($0.6 \leq r < 0.8$) and moderate negative ($-0.8 < r \leq -0.6$) correlations, respectively.

| Module & Pref. stage | Green | Red | Soft | Turning | Black |
|---|---|---|---|---|---|
| 1 (Green) | 0.44 | 0.15 | –0.16 | –0.23 | –0.28 |
| 2 (Red) | –0.17 | 0.52 | –0.06 | –0.23 | –0.13 |
| 3 (Red) | –0.13 | 0.43 | 0.01 | –0.06 | –0.26 |
| 4 (Red) | –0.15 | 0.30 | –0.13 | 0.08 | –0.08 |
| 5 (Red) | 0.09 | 0.28 | 0.14 | 0.00 | –0.50 |
| 6 (Soft) | –0.52 | 0.18 | 0.19 | 0.15 | 0.06 |
| 7 (Turning) | –0.23 | –0.34 | 0.26 | 0.38 | 0.06 |
| 8 (Turning) | 0.15 | –0.29 | 0.26 | 0.29 | –0.31 |
| 9 (Turning) | 0.16 | –0.50 | 0.14 | 0.24 | 0.04 |
| 10 (Black) | –0.06 | –0.25 | –0.18 | –0.02 | 0.50 |
| 11 (Black) | 0.21 | 0.04 | –0.37 | –0.30 | 0.32 |
| 12 (Black) | –0.08 | –0.45 | 0.06 | 0.23 | 0.31 |
| 13 (Black) | –0.41 | –0.20 | 0.15 | 0.24 | 0.30 |
| 14 (Black, Green) | 0.28 | –0.43 | –0.14 | 0.03 | 0.27 |

Correlation coefficient
0.6
-0.6

**Supplementary Figure 12 | Gene regulatory modules inferred by WGCNA.** The heat map illustrates the strength and direction of correlation between module eigengenes and black raspberry fruit ripening stages. For each module, correlation coefficients were calculated by assessing the relationship between transcript abundance patterns and developmental time points, allowing identification of modules associated with specific fruit ripening stages.

A

| Chromosome | Heterochromatin | Euchromatin | Chromosome | Heterochromatin | Euchromatin |
|---|---|---|---|---|---|
| Chr. 1 | 17,550,051 - 21,084,501 | 2,000,001 - 4,750,001 | Chr. 5 | 22,535,801 - 26,022,251 | 2,040,501 - 4,223,151 |
| Chr. 2 | 22,203,451 - 25,285,801 | 5,891,251 - 8,577,201 | Chr. 6 | 46,816,651 - 50,507,801 | 22,753,151 - 26,311,501 |
| Chr. 3 | 16,575,051 - 20,744,351 | 35,637,801 - 39,319,201 | Chr. 7 | 17,415,001 - 22,861,801 | 36,764,651 - 39,821,851 |
| Chr. 4 | 18,013,101 - 21,662,751 | 251,701 - 3,108,901 | | | |

B

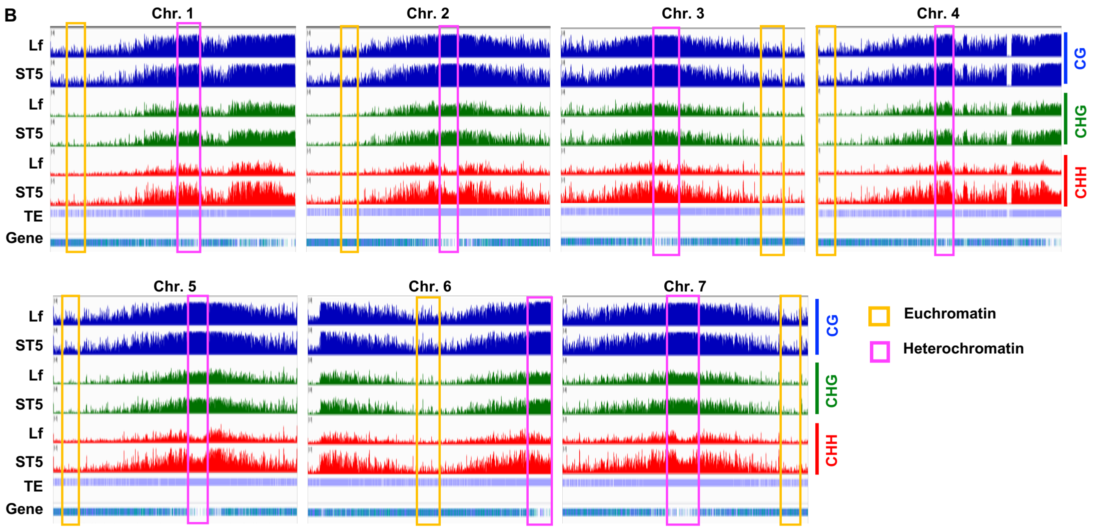


**Supplementary Figure 13 | DNA methylation profiling across seven chromosomes of black raspberry in leaf and black-stage fruit tissues. A** Approximately 2 to 4 Mb genomic intervals from both heterochromatic (transposon-rich) and euchromatic (gene-rich) regions were randomly selected to calculate DNA methylation levels using 50-bp sliding windows, as shown in Fig. 5. **B** Integrative Genomics Viewer of the randomly selected chromosomal segments. Pink and yellow boxes indicate heterochromatic and euchromatic regions, respectively, which were used for finer-resolution methylation analysis based on 50-bp windows. Lf, leaf; TE, transposable element.
